# KCC2 regulates neuronal excitability and hippocampal activity via interaction with Task-3 channels

**DOI:** 10.1101/434571

**Authors:** Marie Goutierre, Sana Al Awabdh, Emeline François, Daniel Gomez-Dominguez, Theano Irinopoulou, Liset Menendez de la Prida, Jean Christophe Poncer

## Abstract

The K^+^/Cl^−^ co-transporter KCC2 *(SLC12A5)* regulates neuronal transmembrane chloride gradients and thereby controls GABA signaling in the brain. KCC2 downregulation is observed in several neurological and psychiatric disorders including epilepsy, neuropathic pain and autism spectrum disorders. Paradoxical, excitatory GABA signaling is usually assumed to contribute to abnormal network activity underlying the pathology. We tested this hypothesis and explored the functional impact of chronic KCC2 downregulation in the rat dentate gyrus. Although the reversal potential of GABAA receptor currents was depolarized in KCC2 knockdown neurons, this shift was fully compensated by depolarization of their resting membrane potential. This effect was due to downregulation of Task-3 leak potassium channels that we show require KCC2 for membrane trafficking. Increased neuronal excitability upon KCC2 suppression altered dentate gyrus rhythmogenesis that could be normalized by chemogenetic hyperpolarization. Our data reveal KCC2 downregulation engages complex synaptic and cellular alterations beyond GABA signaling that concur to perturb network activity, thus offering novel targets for therapeutic intervention.

## Introduction

Fast synaptic inhibition in the brain is primarily mediated by GABAA receptors (GABAAR), which are ligand-gated receptors associated with an anion-permeable conductance. GABAAR currents are carried by fluxes of chloride and, to a lesser extent, bicarbonate ions^1^. Consequently, the polarity of the net ion flux through GABAAR channels relies on both the transmembrane gradients of these ions as well as the neuronal resting membrane potential, which thereby determine the driving force of GABAAR currents. In mature neurons, transmembrane chloride gradients are predominantly regulated by the opposing actions of the two cation-chloride co-transporters (CCCs) KCC2 and NKCC1, which respectively mediate outward and inward co-transport of chloride and potassium ions^2^. Thus, postnatal upregulation of KCC2 expression is associated with a progressive hyperpolarizing shift in the reversal potential of GABAAR-mediated currents (E_GABA_)^3^.

In mature neurons, KCC2 expression and function is rapidly regulated by neuronal activity via multiple posttranslational mechanisms^4,5^. Thus, Ca^2+^ influx through postsynaptic NMDA receptors or during prolonged postsynaptic firing^6^ rapidly reduces KCC2 membrane expression and function through protein phosphatase 1-dependent dephosphorylation of its Ser940 residue and protein cleavage by the calcium-activated protease calpain^7,8,9^. Conversely, chloride influx through GABAA receptors stabilizes KCC2 at the plasma membrane via chloride-mediated inhibition of the serine/threonine WNK1 kinase and its downstream effectors SPAK/OSR1, which phosphorylate KCC2 on Thr906 and Thr1007 residues^10,11,12^. Finally, KCC2 expression is also regulated by several neuromodulators acting on G-protein couples receptors^13^ as well as neurotrophins such as BDNF acting via TrkB signaling^14^.

KCC2 dysregulation has been associated with numerous neurological and psychiatric disorders involving synaptic alterations^15,16^. These include epilepsy^17,18,19,20,21,22,23^, neuropathic pain^24,25^, posttraumatic spasticity^26^, Huntington disease^27^, schizophrenia^28^ and Rett syndrome^29,30,31^. In most cases, reduced KCC2 expression was associated with a depolarizing shift in E_GABA_ that could be partly reversed using NKCC1 antagonists or drugs acting to enhance KCC2 function^18,25,27,30,32^. Since these drugs also ameliorated the pathological symptoms, those were generally assumed to primarily reflect a defect in neuronal chloride transport and altered GABAergic neurotransmission.

However, KCC2 functions in neurons extend beyond the mere control of chloride transport. Through interactions with multiple transmembrane and intracellular partners, KCC2 was shown to regulate dendritic spine morphology^33,34^ and actin cytoskeleton^35,36^, as well as the strength and long-term plasticity of glutamatergic synapses^33,35^. Recent functional proteomics data revealed additional putative KCC2 partners, including some involved in the recycling and trafficking of various transmembrane proteins and receptors^37^. Those may then either influence KCC2 function or, conversely, be regulated by KCC2. Thus, dysregulation of KCC2 expression may affect a variety of neuronal intrinsic and synaptic properties and thereby contribute to abnormal activity patterns that underlie neurological and psychiatric disorders. Deciphering the relative contribution of the various perturbations upon KCC2 downregulation may then help predict the most effective rescue strategies.

In the present study, we therefore explored the functional impact of chronic KCC2 downregulation in the rat dentate gyrus at the cellular, synaptic and network activity levels. We found that KCC2 knockdown had no significant effect on the driving force of GABAAR currents at rest but instead enhanced neuronal excitability and EPSP/spike coupling through membrane depolarization and increased resistance. Pharmacological and biochemical evidence suggest this reflected downregulation of a leak potassium conductance carried by TWIK-related acid-sensitive 3 (Task-3) channels, the expression of which was regulated via interaction with KCC2. KCC2 knockdown was not sufficient to promote epileptiform activity but altered dentate gyrus network activity, which was normalized by restoring granule cell membrane properties. Our results show that KCC2 downregulation affects network activity primarily through enhanced neuronal excitability, suggesting novel targets for therapeutic intervention.

## Results

We investigated the impact of a chronic KCC2 suppression in the dentate gyrus using *in vivo* RNA interference. Young adult rats (P30) were stereotaxically injected in the dorsal dentate gyrus with lentiviruses expressing previously validated KCC2-directed or non-target small hairpin RNA sequences together with GFP^33,35^. Suppression of KCC2 expression in transduced neurons was confirmed by immunofluorescence imaging two weeks after infection (Fig 1A, B). We first analyzed GABAAR-mediated transmission using gramicidin perforated-patch recordings. GABAAR-mediated currents were evoked by somatic laser uncaging of Rubi-GABA (15μM) while varying holding potentials (Fig 1C-D). As expected, the reversal potential of GABAAR currents (E_GABA_) was significantly more depolarized in neurons with suppressed KCC2 expression (−73.26 ± 2.57 mV) compared to control granule cells (−82.10 ± 2.26 mV, t-test, t_24_ = 2.54, p=0.0178, Fig 1E). However, neurons transduced with lentivirus expressing KCC2-directed shRNA also displayed more depolarized resting membrane potential (V_rest_, −83.6 ± 1.52 *vs* −89.77 ± 0.82 mV, t-test, t_24_ = 3.41, p = 0.0023). The shift in E_GABA_ and V_rest_ was of similar magnitude such that the driving force of ion flux through GABAARs at rest was unaffected by KCC2 knockdown (10.34 ± 2.37 *vs* 7.66 ± 2.38 mV, t-test, t_24_ = 0.79, p=0.44). Additionally, neither the amplitude (14.12 ± 1.24 *vs* 14.78 ± 0.72 pA, Mann-Whitney, p=0.42) or mean frequency (0.91 ± 0.13 *vs* 1.17 ± 0.16 Hz, Mann-Whitney, p=0.27) of miniature inhibitory post-synaptic currents (mIPSCs) was affected by chronic KCC2 down-regulation (Fig S1). Therefore, KCC2 suppression in dentate granule cells does not affect steady-state GABAergic transmission, due to an unexpected depolarizing shift in resting membrane potential.

**Figure 1.**
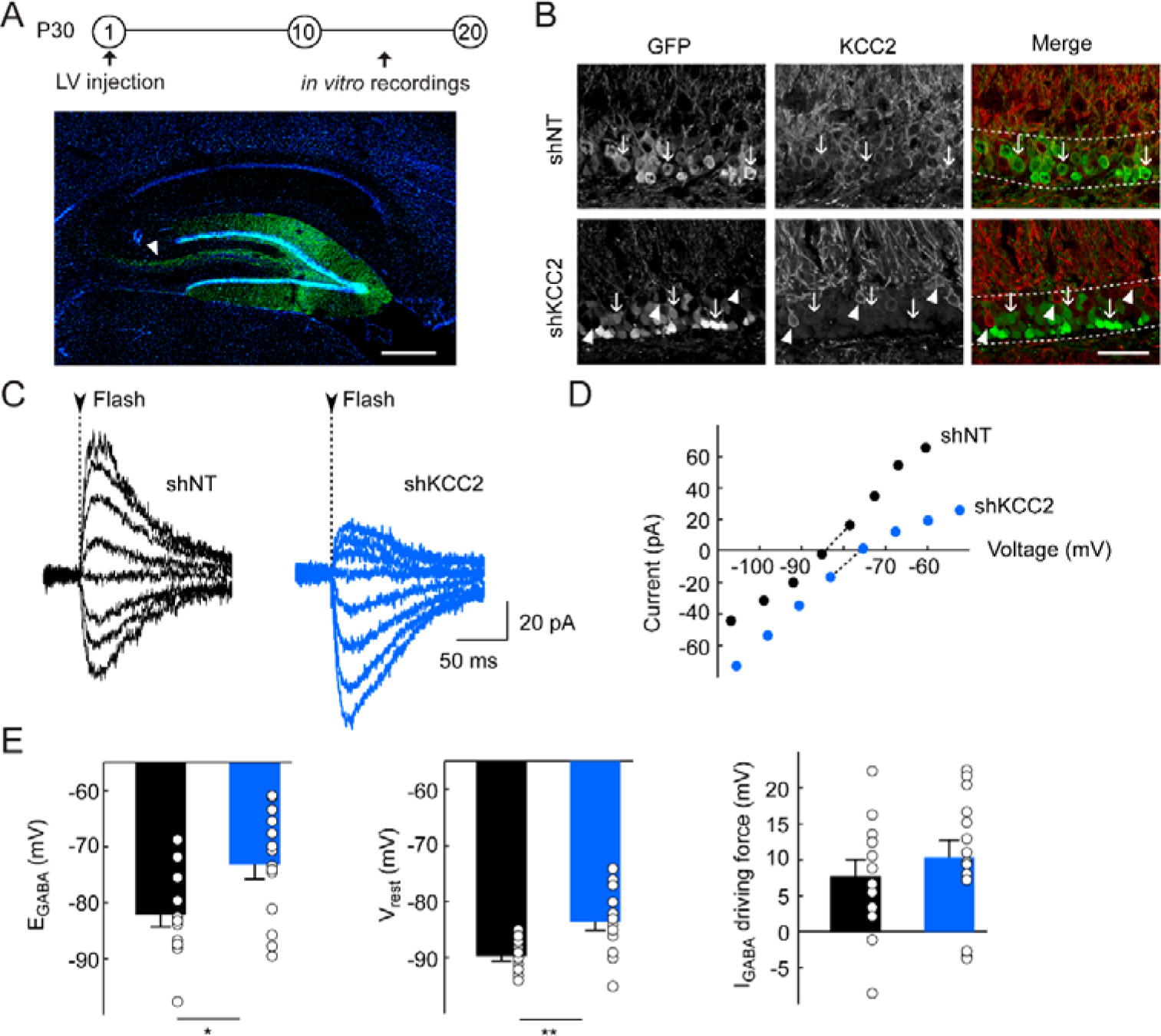
KCC2 suppression in dentate granule cells does not alter steady state GABA signaling. **A**, Top, Experimental timeline. P30 Wistar rats were injected with lentiviral vectors (LV) expressing either non-target (shNT) or KCC2-directed (shKCC2) shRNA sequences and EGFP. *In vitro* electrophysiological recordings were performed 10 to 20 days post-injection. Bottom, maximal intensity projection of confocal micrographs showing infected cells (GFP, green) and DAPI (blue) staining in a parasagittal hippocampal slice 2 weeks after injection. Arrowhead indicates mossy fibers from infected granule cells. Scale, 500 μm. **B**, Maximal intensity projection of confocal micrographs of infected dentate gyrus areas, showing massive extinction of KCC2 expression in KCC2 knockdown neurons. Dotted line delineates *st. granulosum*. Neurons expressing non-target but not KCC2-directed (arrows) show pericellular KCC2 immunostaining, as do neighboring uninfected cells (arrowheads). Scale, 50 μm. **C**, Representative currents following Rubi-GABA somatic uncaging (Flash, dotted line) recorded in gramicidin perforated-patch configuration during incremental voltage steps. **D**, I-V curve of GABA-induced currents corresponding to the examples shown in C. E_GABA_ is interpolated from the two points around reversal. E, Summary data (shNT n = 12 cells, 6 rats, shKCC2 n = 14 cells, 6 rats) show depolarized E_GABA_ upon KCC2 suppression (* p<0.05). The resting membrane potential (V_rest_) is also depolarized (** p<0.01), resulting in no significant change in the driving force of GABAergic currents.

We further explored the mechanisms underlying changes in membrane potential upon KCC2 suppression in granule cells. We recorded infected neurons in whole-cell configuration while blocking synaptic transmission with bicuculline (20 μM), NBQX (20 μM) and APV (50 μM). In these conditions, depolarized resting membrane potential in KCC2 knockdown granule cells (−80.79 ± 1.61 *vs* −87.67 ± 1.06 mV, t-test, t_37_ = 3.45, p = 0.0014) was accompanied by a ~30% increase in input resistance (662.70 ± 47.14 vs 513.38 ± 47.85 MΩ, t-test, t_37_ = 2.21, p = 0.033, Fig 2A). These changes in intrinsic membrane properties resulted in higher excitability of KCC2 knockdown neurons following current injection (Fig 2C, repeated measures ANOVA, F_15,555_ = 11.98, p = 3.18.10^−7^). Changes in the input/output relationship primarily reflected changes of membrane properties since no difference was observed in action potential threshold upon KCC2 knockdown (Fig 2D −50.7 ± 1.92 vs −50.56 ± 1.30 mV, t-test, t_37_ = 0.06, p = 0.95). Additionally, action potential waveform and amplitude (89.6 ± 1.82 vs 88.3 ± 3.31 mV, Mann-Whitney, p = 0.56) were also unchanged in KCC2 knockdown *vs* control neurons (Fig 2E-2F).

**Figure 2.**
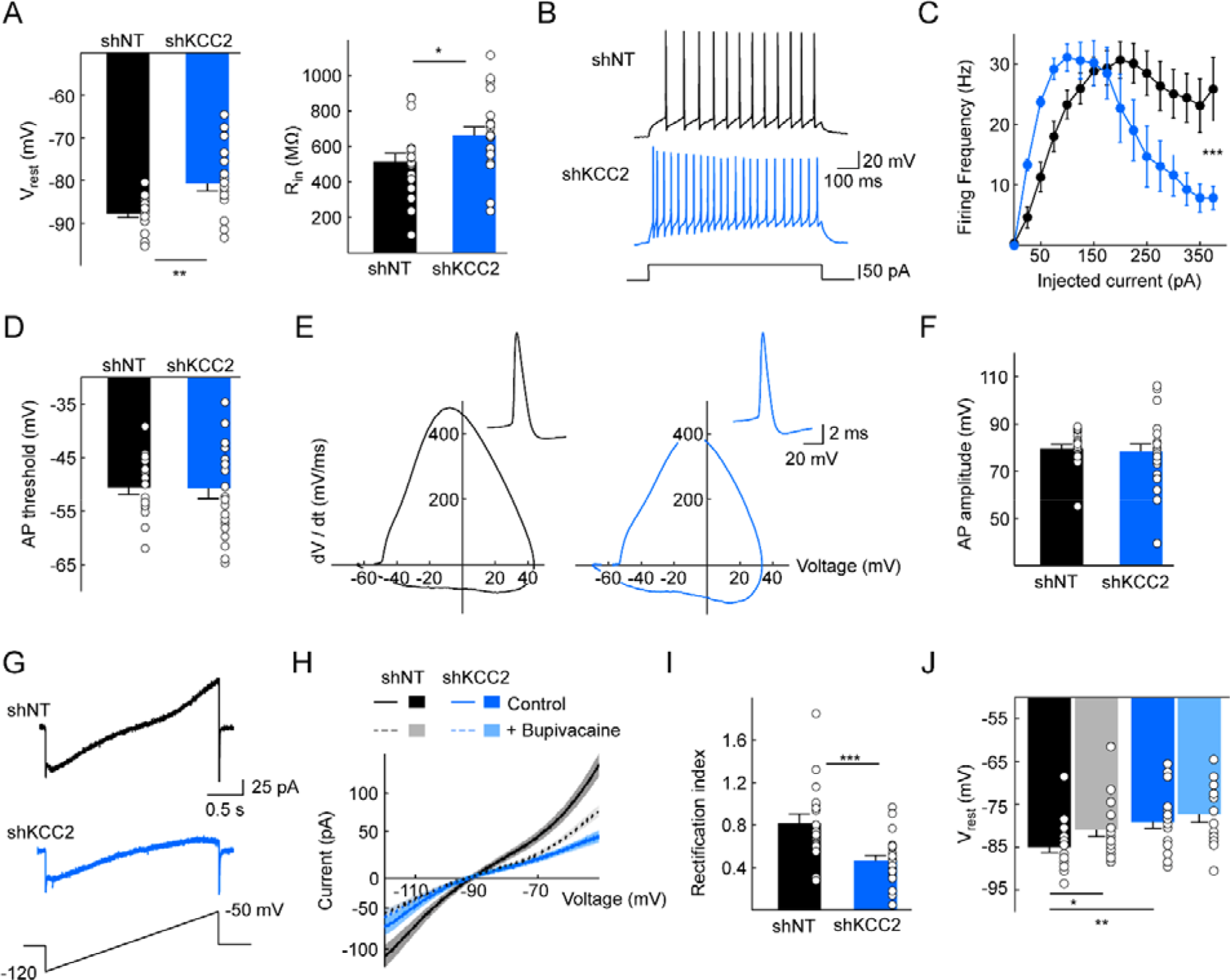
Increased excitability upon KCC2 extinction reflects downregulation of a leak potassium conductance. **A**, Summary graphs of membrane properties of granule cells expressing non-target (n = 18 cells, 5 rats) or KCC2-directed (n = 21 cells, 7 rats) shRNA recorded in whole-cell patch clamp mode in the presence of synaptic transmission blockers. KCC2 knockdown resulted in depolarized membrane potential (** p<0.01) and increased input resistance (* p<0.05). **B**, Individual traces in both conditions for a depolarizing current pulse of 50 pA. **C**, Input/output curve for all cells representing the firing frequency as a function of the injected current (*** p<0.001). **D**, Action potential threshold was determined as the voltage for which derivative of the signal exceeded 15 mV/ms and was not different between conditions. **E**, Individual phase plots of action potentials (insets) recorded in neurons expressing either shRNA sequence, showing no change in the shape of action potentials upon KCC2 suppression. **F**, The amplitude of action potentials was also unchanged in KCC2-knockdown cells. **G**, Representative individual traces for potassium currents recorded in voltage-clamp mode with a voltage ramp ranging −120 to −50 mV at 0.03 mV/ms, in the presence of synaptic transmission blockers and TTX. **H**, Summary plots of mean currents in cells expressing non-target (shNT n = 18 cells, 5 rats) vs. KCC2-directed (shKCC2 n = 21 cells, 7 rats) shRNA. Bath application of the leak-potassium channel blocker bupivacaine (100 μM) reduced potassium currents only in control cells (shNT n = 16 cells, 3 rats, shKCC2 n = 19 cells, 5 rats). **I**, Rectification index plot, defined as the ratio of potassium-mediated currents at −60 mV vs. −120 mV. KCC2 knockdown resulted in reduced rectification (*** p<0.001). **J**, The difference in V_rest_ between control and KCC2-knockdown neurons is almost entirely suppressed by bath application of bupivacaine (100μM) (* p<0.05).

Increased neuronal excitability upon KCC2 down-regulation was cell-autonomous since neighboring uninfected cells had similar properties in slices from animals infected with viruses expressing non-target or KCC2-directed shRNA (Fig S2). In addition, acute application of the KCC2-specific antagonist VU0463271^38^ (6 μM) for more than 30 min did not affect membrane resistance in granule cells from uninfected animals (298.95 ± 26.16 vs 356.14 ± 35.73 MΩ, t_30_ = 1.31, p = 0.20, Fig S2), indicating that KCC2-mediated ion transport *per se* does not significantly contribute to input resistance. Finally, these modifications were not specific of dentate granule cells, since KCC2 knockdown in CA1 pyramidal cells also resulted in depolarized resting membrane potential (−66.59 ± 1.04 vs −72.06 ± 1.53 mV, t-test, t_20_ = 2.95, p = 0.0079), increased input resistance (213 ± 25.38 vs 134.45 ± 10.21 MΩ, Mann-Whitney, p = 0.0013) and shifted input/output curve (repeated measures ANOVA, F_15,300_ = 11.5, p = 0.0029, Fig S3).

Depolarized membrane potential and increased input resistance likely reflect a reduction in potassium currents operating at rest. We tested this hypothesis by recording potassium currents upon voltage ramps from −120 to −50 mV at 0.03 mV/ms while blocking synaptic transmission. In KCC2 knockdown granule cells, potassium conductance was reduced throughout the range of potentials tested (Fig 2G-H). This reduction however was of higher magnitude for outward than inward currents, as evidenced by reduced rectification index (I_−60_/ I_−120_) in KCC2 knockdown *vs* control granule cells (0.46 ±0.05 vs 0.82 ± 0.08, t-test, t_37_ = 3.62, p = 0.0009, Fig 2I). This suggests that outwardly rectifying leak potassium channels might be down-regulated upon KCC2 suppression. Two-pore-domain potassium channels (K2P) are a large family of leak potassium channels comprising both inward and outward rectifiers^39,40^. We therefore tested whether currents carried by K2P channels may be reduced in KCC2 knockdown neurons. Bath application of bupivacaine (100μM), a non-selective K2P channel antagonist significantly reduced potassium currents in neurons expressing non-target shRNA but had very little effect in neurons expressing KCC2-directed shRNA (Fig 2G). Moreover, bupivacaine abolished the difference in resting membrane potential (−77.31 ± 1.86 vs −81.03 ± 1.48 mV, Mann-Whitney test with Bonferroni correction, p = 1, Fig 2J) and input resistance (998.7 ± 118.6 vs 1343.1 ± 207.4 MΩ, Mann-Whitney test with Bonferroni correction, p = 1) between KCC2 knockdown and control granule cells. Together, these results demonstrate that chronic KCC2 suppression increases neuronal excitability primarily through regulation of an outwardly rectifying leak potassium conductance.

Trek-2 and Task-3 are the most prominent outward-rectifier K2P in dentate gyrus granule cells^41,42,43^. Immunohistofluorescence imaging of Trek-2, however, revealed a predominant expression in mossy fibers (Fig S4A) with little or no expression in somato-dendritic compartments, suggesting Trek-2 may not prominently contribute to granule cell resting membrane potential and membrane conductance. Conversely, Task-3 expression was primarily observed in granule cell somata and dendrites (Fig S4B). We therefore examined whether Task-3 protein expression in granule cells might be affected by chronic KCC2 suppression. Immunofluorescence confocal imaging indeed revealed a reduced expression of Task-3 in granule cells expressing KCC2-directed shRNA compared to non-target shRNA (Fig 3A). We then investigated the mechanisms underlying Task-3 down-regulation upon KCC2 silencing. KCC2 has been shown to interact with a variety of proteins, including transmembrane proteins such as GluK2^44^, Neto2^45^ and GABA_B_ receptors^46^ that regulate KCC2 membrane expression or the intracellular guanylyl exchange factor βPIX which is regulated through direct interaction with KCC2^35,36^. We therefore asked whether KCC2 might also interact with Task-3 channels and thereby influence their membrane expression and/or function. In order to test whether KCC2 interacts with Task-3 channels in neurons, we performed a co-immunoprecipitation assay from adult rat hippocampal homogenates. We observed that anti-Task3 antibodies pulled down KCC2, indicating an interaction between the endogenous proteins in hippocampal neurons (Fig 3B). In order to characterize KCC2-Task-3 interaction further, we co-immunoprecipitated exogenously expressed Flag-tagged KCC2 and HA-tagged Task-3 in HEK293T cells. In lysates from double-transfected cells, anti-HA antibodies pulled down Flag-KCC2 together with HA-Task-3 while anti-Flag antibodies pulled down HA-Task-3 and Flag-KCC2 (Fig 3C). Importantly, this interaction was specific for Task-3 since overexpressed HA-tagged GFP failed to interact with KCC2 in a similar assay (Fig. 3C). We then asked whether interaction with KCC2 may influence the expression of Task-3 channels. To test this hypothesis, we expressed HA-Task-3 in Neuro2A cells either alone or together with Flag-KCC2 and analyzed Task-3 expression by immunostaining and confocal imaging. In the absence of KCC2, Task-3 expression at plasma membrane was reduced and accumulated in an intracellular compartment (Fig. 3D-E). Line scans of immunofluorescence across the cell membrane and cytoplasm therefore revealed a reduced membrane to intracellular intensity ratio in cells lacking KCC2 (0.74 ± 0.08 vs. 2.47 ± 0.34, Mann-Whitney, p = 1.63.10^−7^, Fig 3E). Together, our results show that KCC2 interacts with Task-3 and regulates its membrane expression in dentate gyrus granule cells, thereby regulating their intrinsic excitability.

**Figure 3.**
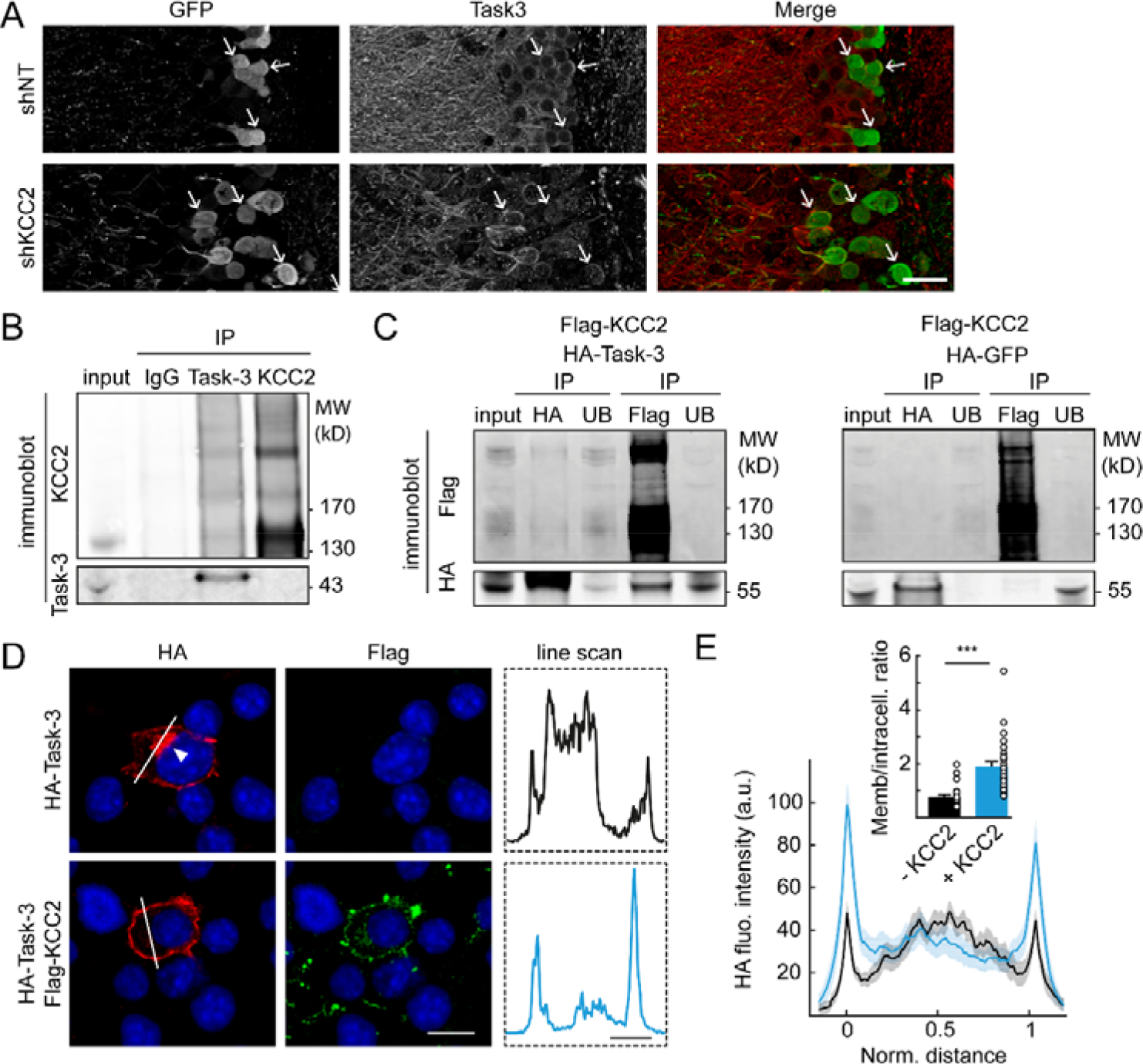
KCC2 interacts with Task-3 and promotes its membrane expression. **A**, Maximal intensity projection of confocal micrographs (4 focal planes, 0.2 μm z-step) of infected dentate gyrus, showing decreased Task-3 pericellular immunostaining in KCC2 knockdown *vs.* control granule cells. Scale, 10 μm. **B**, Immunoblot of co-IP samples of endogenous proteins extracts from adult rat hippocampi. Solubilized membrane homogenates (input) were immunoprecipitated with either Task-3, KCC2 or control IgG antibodies. Eluates resulting from immunoprecipitations (IP fractions) were then probed for both Task-3 and KCC2. Task-3 IP pulled down KCC2, therefore demonstrating KCC2 interaction with Task-3 in adult hippocampus. KCC2 IP however did not reveal detectable Task-3 protein level in these samples. Representative example of three independent replicates. **C**, Immunoblot of the immunoprecipitates from HEK293T cell homogenates coexpressing recombinant, Flag-tagged KCC2 and HA-tagged GFP (left) or HA-tagged Task-3 (right). Immunoprecipitation using Flag antibodies pulled down Task-3 (right) but not GFP (left). Conversely, immunoprecipitation with HA antibodies precipitated Flag-KCC2 only in HA-Task-3 expressing cells. Each immunoblot is a representative example of three independent experiments. **D**, Focal plane from confocal micrographs of HEK293T cells transfected with either HA-Task-3 alone (top) or HA-Task-3 together with Flag-KCC2 (bottom). Note the accumulation of HA immunostaining in an intracellular compartment in absence of Flag-KCC2 (arrowhead). Scale, 20μm. Fluorescence intensity of HA staining across the cell membrane and cytoplasm was analyzed along the white line shown on the micrographs and plotted in dotted insets (right). Scale, 5 μm. **E**, Averaged, scaled line scan intensity plots from all cells (HA-Task-3, n = 24; HA-Task-3+Flag-KCC2, n=27; 3 independent cultures) reveal intracellular accumulation of Task-3 in absence of KCC2 as evidenced by the averaged HA intensity across line scans and the decreased ratio of membrane to intracellular intensity (inset, *** p<0.001).

We next asked how reduced leak conductance and enhanced excitability upon KCC2 suppression might affect EPSP/spike coupling in granule cells. Previous work showed that loss of KCC2 expression in hippocampal neurons led to reduced synaptic strength at glutamatergic inputs due to enhanced lateral diffusion of AMPA receptors^33^. Enhanced intrinsic excitability may then act as a homeostatic compensatory mechanism for reduced synaptic excitation^47^. To test this hypothesis, we first compared synaptic strength at glutamatergic inputs onto granule cells by recording miniature excitatory postsynaptic currents (mEPSCs). As previously reported, KCC2 knockdown granule cells showed a ~20% decrease in mEPSC amplitude (5.06 ± 0.43 vs 6.54 ± 0.28 pA, Mann-Whitney, p = 0.0009, Fig 4A-B) with no change in their mean frequency (1.60 ± 0.42 vs 1.82 ± 0.41 Hz, Mann-Whitney, p = 0.09, Fig 4A-B) compared to control cells. From these recordings, waveforms of quantal currents were derived, fit and used as a current command in current-clamp recordings from KCC2 knockdown *vs* control granule cells. Thus, neurons were recorded in whole-cell configuration and maintained at their resting membrane potential while somatically injecting multiples of quantal current waveforms to mimic synchronous excitatory inputs (Fig 4C). Although the efficacy of excitatory inputs was reduced in KCC2 knockdown neurons compared to control, enhanced intrinsic excitability overcompensated this reduction, leading to enhanced EPSP/spike coupling. Thus, KCC2 knockdown neurons fired action potentials from ~40 simultaneous quanta whereas firing of control neurons expressing non-target shRNA required at least 60 quanta (Fig 4C, repeated measures ANOVA, F_20,500_ = 4.83, p = 0.0374). In addition, synaptic integration during high frequency (>50 Hz) trains of EPSCs was also facilitated in KCC2 knockdown granule cells, owing to increased membrane resistance compared to control cells (Fig. 4D).

**Figure 4.**
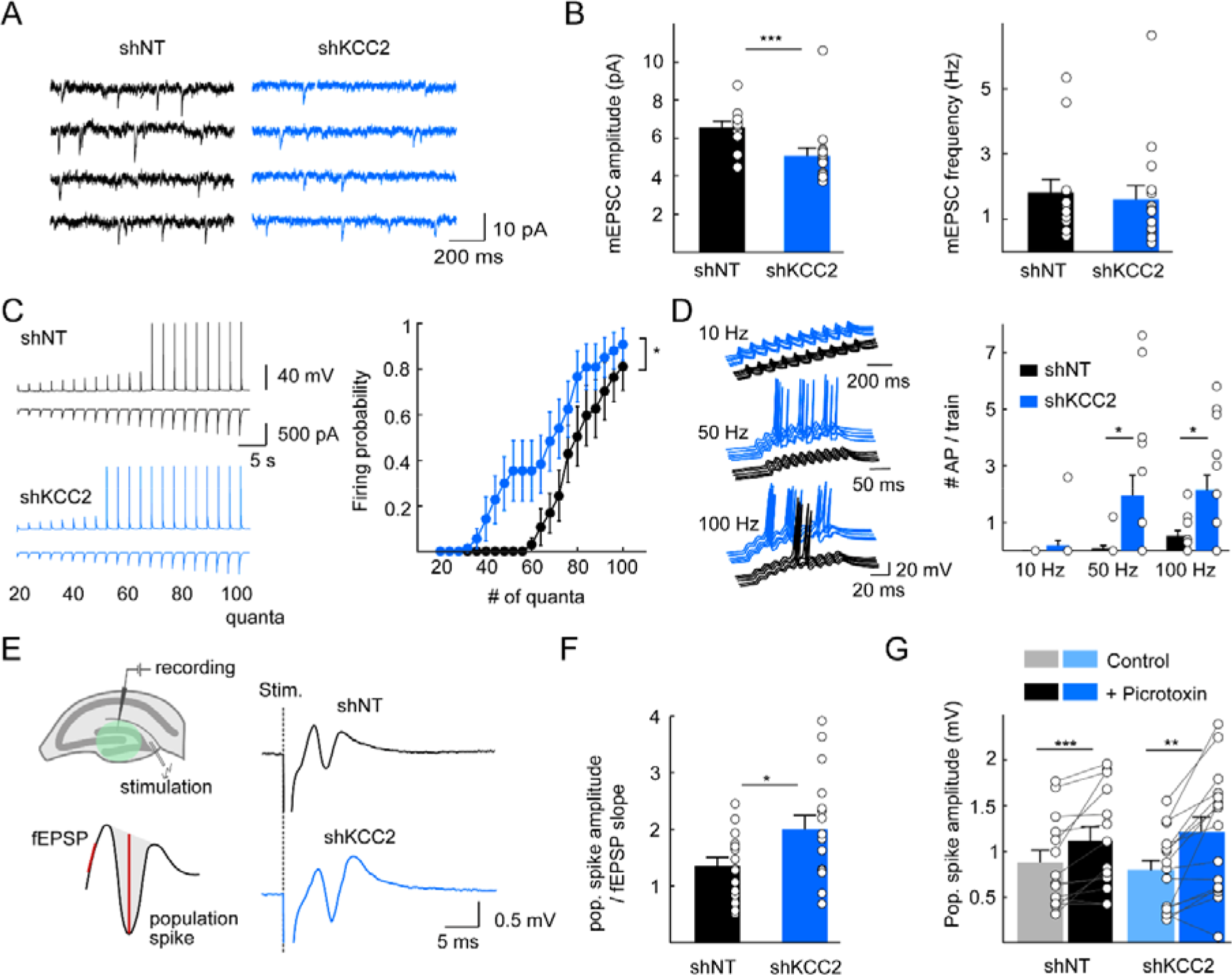
KCC2 suppression increases EPSP/Spike coupling in dentate granule cells. **A**, 4 s-recordings of mEPSCs from infected granule cells expressing non-target (shNT, black, n = 13 cells, 3 rats) or KCC2-directed (shKCC2, blue, n = 15 cells, 3 rats) shRNA. **B**, Summary data of mEPSC showing reduced amplitude (left) but not frequency (right). **C**, Left, representative recordings of current clamp recordings (top) from granule cells kept at their resting membrane potential upon somatic injection of multiples of scaled, mEPSC waveforms of increasing amplitude (bottom). Right, summary plots of all data (shNT, n = 13 cells, 2 rats, shKCC2, n = 15 cells, 2 rats) showing enhanced EPSP/spike coupling in KCC2 knockdown neurons (* p < 0.05). **D**, Neurons were kept at −87 mV in current clamp mode and trains of 120 pA amplitude mEPSCs (about 20x quantal size) were injected as in C at various frequency. Left, representative traces for trains of 10 EPSC waveform at 10, 50 and 100 Hz. Right, for trains delivered at 50 or 100 Hz, firing probability was enhanced in KCC2 knockdown granule cells (* p< 0.05, shNT, n = 13 cells, 2 rats, shKCC2, n = 15 cells, 2 rats). **F**, Left, experimental paradigm to test EPSP/spike coupling for synaptically evoked responses in slices. A bipolar electrode was located in the molecular layer to stimulate entorhinal afferents while field PSPs and population spikes were recorded with a glass electrode located in the granule cell layer (top). fEPSP and population spike amplitudes were measured as shown (bottom). Right, average of 10 consecutive responses recorded at 0.1 Hz. **G**, Data from n = 15 slices infected with non-target shRNA (shNT, 6 animals,) and 16 slices infected with KCC2-directed shRNA (shKCC2, 6 animals) confirmed increased coupling of fEPSP to population spike (* p<0.05). **H**, Bath application of the GABAAR pore blocker picrotoxin resulted in increased amplitude of the population spike in both conditions, indicating GABA transmission was functionally inhibitory.

Perforant path input to the dentate gyrus provides synaptic excitation to granule cells as well as feedforward inhibition that contributes to their sparse activation by entorhinal afferents^48^. In order to explore EPSP/spike coupling of perforant path inputs in a more physiological setting, we recorded evoked fEPSP and population spike upon stimulation of perforant path inputs. The recording electrode was positioned in a densely infected area of the granular layer and stimulus intensity was set such that the population spike amplitude was about half its maximum amplitude (see Methods). In slices infected with lentiviruses expressing KCC2-directed shRNA, fEPSP/population spike coupling was increased by ~47% compared to control slices expressing non-target shRNA (2.00 ± 0.25 vs 1.36 ± 0.15 ms^−1^, t-test, t_29_ = 2.18, p = 0.0374, Fig 4F). This result further supports that increased excitability upon KCC2 knockdown is not homeostatic but instead acts to promote granule cell recruitment by entorhinal afferents. Importantly, bath application of the GABAAR antagonist picrotoxin (100μM) increased the amplitude of population spike similarly in slices expressing either non-target shRNA (1.12 ± 0.15 mV *vs* 0.88 ± 0.13 mV, paired t-test, t_13_ = 3.33, p = 0.0054) or KCC2-directed shRNA (1.21 ± 0.17 mV *vs* 0.79 ± 0.11 mV, paired t-test, t_15_ = 4.29, p = 0.0006; Fig 4G). These results suggest that GABAergic transmission, although depolarizing in dentate granule cells^49^ (Fig. 1E), remains shunting and inhibitory, independent of KCC2 expression. Therefore, KCC2 suppression in dentate granule cells promotes their recruitment by entorhinal afferents primarily through enhanced excitability and EPSP/spike coupling rather than altered GABA signaling.

The dentate gyrus is often considered as a filter or a gate for activity propagation from the entorhinal cortex to the hippocampus^50^, in particular owing to dentate granule cells’ sparse firing. These properties allow for pattern separation^51^ and may prevent runaway excitation in the hippocampus^52^. Enhanced EPSP/spike coupling in dentate granule cells upon KCC2 suppression may then act to increase their excitatory drive by entorhinal afferents and thereby alter hippocampal rhythmogenesis. We explored this hypothesis using chronic intrahippocampal recordings from rats injected with lentivirus expressing either KCC2-directed or non-target shRNA (n=7 and 5, respectively, Fig 5A-B). In these animals, about 40-70% granule cells were infected over 1-2 mm around the injection site (Fig. 5B and Fig S5). Unlike a recent study using a similar approach^53^, we did not observe spontaneous seizures in any of these animals. Susceptibility to pilocarpine-induced seizures was also unaffected by KCC2 knockdown in the dorsal dentate gyrus (Fig S6A-C). Chronic, intrahippocampal recordings using linear silicon probes did not reveal interictal spikes or fast ripples (Fig S6D-E) that could represent hallmarks of an epileptic hippocampal network even in the absence of behavioral symptoms^54^. These results indicate that focal KCC2 suppression in the dentate gyrus is not sufficient to trigger or promote epileptiform activity.

**Figure 5.**
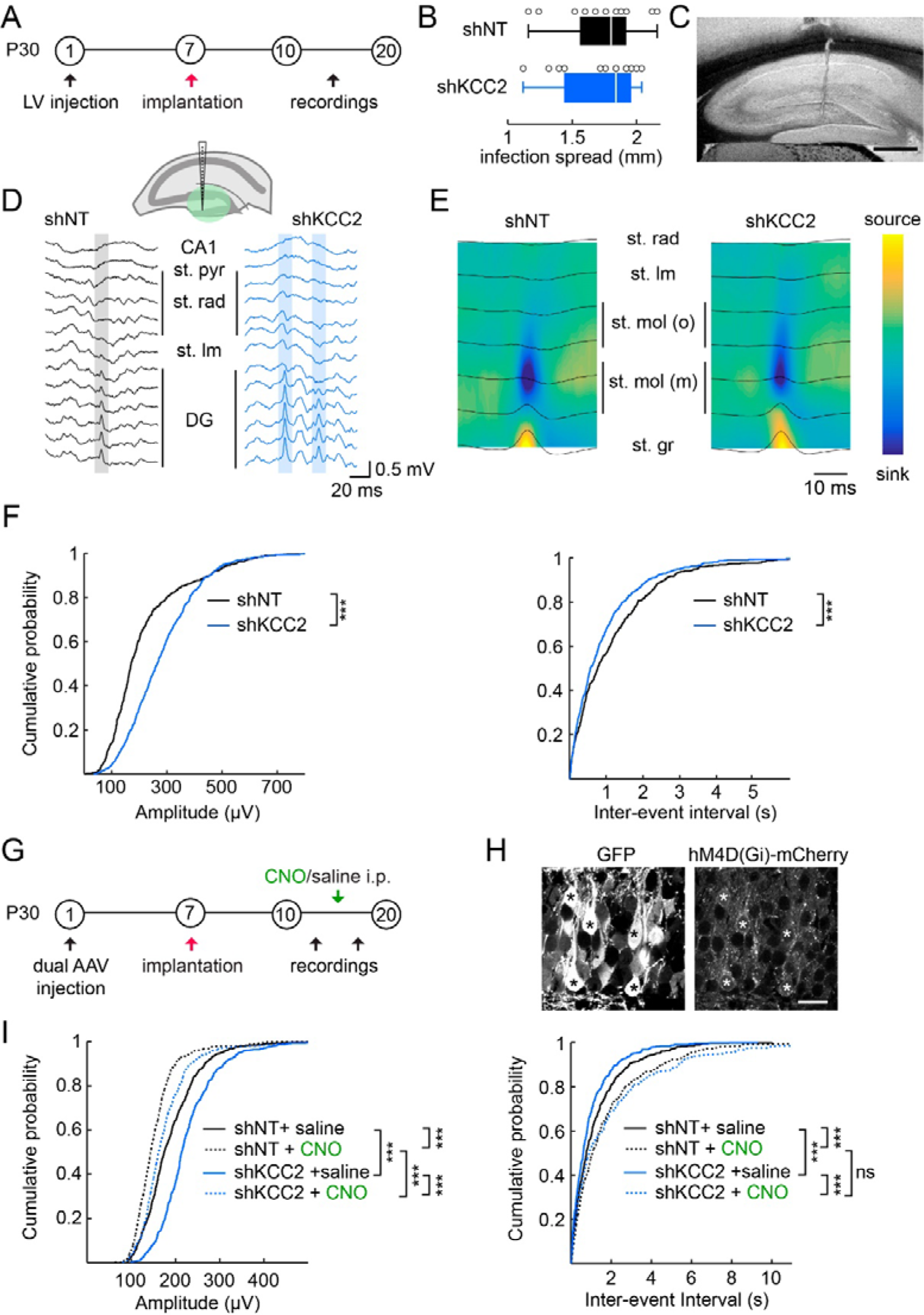
KCC2 suppression alters dentate gyrus activity primarily through increased neuronal excitability. **A**, Experimental timeline. P30 Wistar rats were injected with shRNA-expressing lentiviral vectors. A week later, 16-channels linear silicon probe were implanted at the same coordinates. *In vivo* recordings were performed starting from 3 days post-implantation. **B**, Longitudinal spread of the viral infection was determined at the end of each experiment and was similar between groups (shNT n= 5 rats, shKCC2 n = 7 rats; p= 0.45). **C**, Macroscope DIC micrograph revealing probe localization (vertical scar). Scale, 250 μm. **D**, Top, the 16-channels silicon probe allow simultaneous recording of all hippocampal layers. Examples of recordings during non-REM sleep showing dentate spikes (highlighted in grey: shNT or blue: shKCC2). **E**, Dentate spikes were automatically detected for each animal. Averaged current source density plots over 100 events per animal reveal similar profiles between groups and allow identification of granule cell layer (presence of a source). **F**, Dentate spikes were automatically detected for each animal. Cumulative amplitude (left) and inter-event intervals (right) histograms from 140 randomly selected events per animal, showing increased amplitude and frequency of dentate spike upon KCC2 knockdown in the dentate gyrus (*** p<0.001) **F**, Experimental timeline for rescue experiments with DREADD silencing as above, except that each animal was after either saline or CNO (1 mg/kg) i.p. injection. **G**, Single plane confocal micrographs confirmed most granule cells were co-infected with shRNA+GFP and hM4D(Gi)-mCherry (stars). Scale, 20 μm. **H**, Dentate spikes were automatically detected for each animal (shNT n = 4 rats, shKCC2 n = 3 rats). Cumulative amplitude (left) and inter-event intervals (right) histograms from 185 randomly selected events per animal. Injection of CNO significantly reduced dentate spike amplitude (p < 0.001) and frequency (p < 0.001) in both groups of animals.

Further analysis of hippocampal rhythmogenesis during various behavioral states however revealed significant alterations upon KCC2 knockdown in the dentate gyrus. Thus, while theta-band activity recorded during exploration and rapid-eye movement (REM) sleep was not significantly affected (Fig S7), we observed a specific increase in both the amplitude (Kolmogorov-Smirnov, p < 0.001, Fig 5F) and frequency (Kolmogorov-Smirnov, p < 0.001, Fig 5E) of dentate spikes during slow-wave sleep (SWS). Dentate spikes are sharp hilar LFP events associated with rest^55^ that arise from a transient increase in the excitatory drive from the entorhinal cortex, as revealed by current source density analysis^55^ (Fig. 5D-E). Our results suggest these changes in dentate spike activity resulted primarily from increased excitability of dentate granule cells. Restoring granule cell excitability independently of chloride export and GABA signaling was then predicted to rescue dentate spike activity. We tested this prediction by co-injecting animals with adeno-associated viruses (AAV2.1) expressing the inhibitory receptor hM4D(Gi)^56^ and either non-target or KCC2-directed shRNA with GFP. Using this approach, all granule cells expressing GFP also expressed hM4D(Gi), thereby enabling selective silencing of neurons expressing shRNA sequences (Fig. 5G-H). We first verified that hM4D(Gi) activation was sufficient to hyperpolarize transduced granule cells *in vitro*. Indeed, in whole cell recordings from transduced neurons, bath application of the hM4D(Gi) ligand CNO (10μM) induced a hyperpolarizing shift of V_rest_ by −5.66 ± 1.42 mV (paired t-test, t_5_ = 4.36, p = 0.0073, Fig S8). As in lentivirus-infected animals, AAV-mediated KCC2 suppression in the dorsal dentate gyrus increased the amplitude and reduces the frequency of dentate spikes during SWS (n = 4 rats infected with non-target shRNA and 3 with KCC2-directed shRNA, Kolmogorov-Smirnov, p < 0.001, Fig 5I). Following systemic injection of CNO (1 mg/kg, i.p.), the amplitude of dentate spikes in animals injected with KCC2-directed shRNA was reduced to levels below those of saline-treated control animals (Fig 5I). Similarly, CNO injection reduced the frequency of dentate spikes in both KCC2-knockdown and control animals (Kolmogorov-Smirnov, p < 0.001, Fig 5I). Together, our data demonstrate that KCC2 suppression in dentate granule cells leads to altered dentate gyrus activity through a reduced Task3-mediated potassium conductance and a subsequent increase in neuronal excitability, independent of GABA signaling.

## Discussion

Cation-chloride cotransporters (CCCs) are essential to neuronal chloride homeostasis. Thus, changes in their expression level in the developing brain are associated with changes in the polarity of GABA transmission^3^ and altered CCC function or expression is observed in a variety of neurological and psychiatric conditions^15,16^. In the pathology, KCC2 downregulation is often assumed to primarily affect GABA signaling and thereby promote hyperexcitability^17,18,25,26,27,57^. However, this hypothesis was not tested in isolation from other, potentially confounding pathological factors. Here we directly tested the impact of a chronic KCC2 suppression both on the cellular and synaptic properties of hippocampal neurons as well as on network activity associated with behavior. We demonstrate that KCC2 knockdown in dentate granule cells has little effect on GABA signaling at rest but instead increases their excitability via interaction with the leak potassium channel Task-3. We establish that this increased excitability strengthens EPSP/spike coupling in dentate gyrus granule cells thereby acting to enhance their recruitment by entorhinal afferents. This in turn results in altered local rhythmogenesis that was normalized by restoring granule cell intrinsic excitability. Thus, KCC2 downregulation affects a variety of synaptic and intrinsic properties in hippocampal neurons beyond the mere control of GABA transmission and concur to perturb hippocampal activity.

KCC2 interacts with a variety of transmembrane as well as intracellular partners, including postsynaptic receptors^44,45,46^, actin-related proteins^34,35,36^ and others involved in protein trafficking and recycling^37^. Although these interactions are most often considered with regard to KCC2 expression or function, they also influence the function of KCC2 partners. For instance, loss of βPIX-KCC2 interaction increases the activity of βPIX effectors of the Rac1/PAK/LIMK pathway, likely through subcellular redistribution^35,36^. Here, using immunoprecipitation assays, we identified the leak potassium channel Task-3 (*KCNK9*) as a novel KCC2 interactor in hippocampal neurons. We show Task-3 membrane expression and function in neurons are reduced upon KCC2 downregulation and demonstrate that Task-3 membrane targeting requires KCC2 expression in heterologous cells. Thus, in cells lacking KCC2, Task-3 was sequestered intracellularly and its membrane expression was reduced (Fig. 3). We did not attempt to fully identify the molecular determinants of this interaction or the mechanism involved in KCC2-dependent trafficking of Task-3 channels. However, it is remarkable that whereas Task-3 was not found among KCC2 interactors by functional proteomics, several members of the *YWHA* family encoding 14-3-3 proteins were identified as putative partners^37^. 14-3-3 are adaptor proteins involved in a variety of biological functions that include masking phosphorylated or interaction sites of their targets or altering their subcellular localization^58^. In particular, 14-3-3 proteins were shown to bind to and influence the membrane traffic of both Task-1 and Task-3 channels^59^. This effect is mediated through a KRR retention signal in the carboxy-terminal domain of the channels that is masked by 14-3-3 binding. Thus, phosphorylation-dependent binding of 14-3-3 suppresses coatomer protein I (COPI)-mediated retention of Task channels in the endoplasmic reticulum^59,60^. We therefore propose that KCC2 binding to both 14-3-3 and Task-3 may somehow influence the interaction between the two proteins, thereby preventing COPI-mediated intracellular sequestration of the channels. We cannot exclude, however, that KCC2-Task-3 interaction may also occur at the plasma membrane and act to stabilize Task-3 channels.

In mature neurons, KCC2 plays a prominent role in maintaining transmembrane chloride gradients, thereby influencing ion flux through the chloride-permeable GABAA receptor channels. Thus, KCC2 pharmacological blockade^45,46^, genetic ablation^61,62^ or suppression^63^, or activity-dependent modulation^6,11,46,64^ all result in a depolarization of the reversal potential of GABAAR-mediated currents (E_GABA_). Consistent with these observations, we found that chronic KCC2 knockdown in dentate granule cells resulted in a depolarizing shift of E_GABA_ by about 9 mV. This effect was, however, almost fully compensated by a depolarization of their resting membrane potential (V_rest_) of about the same magnitude (Fig. 1), resulting in a driving force of GABAAR currents that was virtually unaffected at rest. This effect was not specific of dentate gyrus granule cells as it was observed also in CA1 pyramidal neurons (Fig. S3). Despite the importance of the membrane potential in setting the efficacy and polarity of GABA signaling, only one study investigated the effect of KCC2 suppression on V_rest_ in cerebellar neurons^61^. Thus, conditional KCC2 ablation resulted in a depolarization of V_rest_ that also compensated the shift in E_GABA_ in granule cells but not Purkinje cells. Interestingly, hippocampal and cerebellar granule cells as well as CA1 pyramidal cells strongly express Task-3 channels^42,65^ whereas in Purkinje cells predominantly express Task-1 and Task-565,66. Thus, differential expression of leak potassium channels may underlie cell-specific effects of KCC2 knockdown on resting membrane potential and the driving force of GABAAR currents.

KCC2 downregulation has often been linked to epileptogenesis. Thus, KCC2 expression is reduced in several animal models of experimental epilepsy including kindling^67^, traumatic brain injury^68^ and pilocarpine-induced *status epilepticus^20^*. Moreover, genetic studies identified loss of function mutations in the *Slc12a5* gene encoding KCC2 in families with idiopathic epilepsy^19,22,69^. Finally, lack of KCC2 expression was associated with depolarized E_GABA_ in 20-30% principal neurons in the subiculum resected from temporal lobe epilepsy patients^18,70^ as well as in human peritumoral neocortical slices^71^. This work suggested that KCC2 downregulation may be causal to the pathological, interictal activity recorded *in vitro* and a modeling study supported such conclusion^17^. Our experiments allowed us to test this hypothesis experimentally. Despite a high density and widespread infection in most of the dorsal dentate gyrus, we did not however find any sign of epileptiform activity in rats with suppressed KCC2 expression (Fig. S5). Thus, these animals did not show spontaneous seizures and intrahippocampal recordings around the infected area failed to reveal interictal discharges or fast ripples that occur between seizures both in patients and in animal models of focal epilepsy^72,73^. Susceptibility to pilocarpine-evoked *status epilepticus* was also unaffected by KCC2 knockdown. Our data therefore argue against a causal link between KCC2 downregulation and the emergence of epileptiform activity, at least in the dentate gyrus. Several arguments may help resolve this apparent discrepancy between our results and data from human epileptic tissue and animal models. First, epileptic networks undergo a number of physio-pathological alterations that may contribute to or promote epileptiform activities, including cell death, axonal sprouting, inflammation, as well as changes in synaptic and intrinsic neuronal membrane properties^74^. Thus, it is interesting to note that KCC2 suppression failed to produce epileptiform activity in a subicular model network unless extracellular potassium was raised to a non-physiological range, thereby increasing neuronal excitability^17^. In addition, our data show that depolarized E_GABA_ upon KCC2 suppression is fully compensated by depolarized V_rest_, such that KCC2 knockdown neurons are more excitable although GABA signaling remains inhibitory. Depolarizing GABAergic responses may then require further depolarization of E_GABA_, for instance through concomitant upregulation of the NKCC1 transporter, as observed in temporal lobe epilepsy models^20,75^ as well as in the cortex and hippocampus from epileptic patients^71,76^.

Our results show that KCC2 downregulation in the dentate gyrus is nevertheless sufficient to alter local rhythmogenesis. Thus, we report a specific increase in the amplitude and frequency of dentate spikes upon KCC2 knockdown (Fig. 5). Dentate spikes are population activities that occur in the dentate gyrus during sleep and immobility and are thought to represent synchronous activation of granule cells by entorhinal afferents^55^. This phenotype is therefore consistent with the cellular impact of KCC2 knockdown we observed in dentate granule cells, i.e. membrane depolarization and increased excitability due to Task-3 downregulation. We showed these effects combined to promote EPSP/spike coupling, despite a reduced synaptic strength at glutamatergic inputs onto granule cells, as previously observed in hippocampal neurons^33,35^. Although granule cells fire sparsely under physiological conditions, KCC2 suppression and subsequent increased excitability promote their recruitment by entorhinal afferents. Thus, fewer simultaneous quantal EPSCs are required for firing KCC2 knockdown granule cells and they fire more action potentials than control cells during high frequency trains of afferent inputs (Fig. 4C-D). Several arguments support that these effects were mostly independent of KCC2-induced alterations in GABA transmission. First, GABA transmission was still inhibitory upon KCC2 suppression, as indicated by the unchanged driving force of GABAAR-mediated currents (Fig. 1) and the excitatory effect of picrotoxin on evoked population spikes (Fig. 4G). Second, increased dentate spike amplitude was fully compensated by chemogenetic hyperpolarization of KCC2 knockdown neurons (Fig. 5). Since GABAAR activation is depolarizing in dentate granule cells^49^, hM4D(Gi)-induced hyperpolarization rescued V_rest_ in infected granule cells but further enhanced the depolarizing nature of GABA transmission. If GABA signaling was involved in the increased dentate spike amplitude and frequency, hM4D(Gi) silencing would then be expected to worsen, not rescue this phenotype.

Together, our work reveals for the first time the effects of a chronic downregulation of KCC2 expression in an intact network. We show these effects are diverse but primarily involve increased neuronal excitability, not reversed GABA signaling. They lead to altered network activity without however promoting seizures. Thus, KCC2 suppression in the dentate gyrus primarily affects dentate spike activity with no detectable effect on theta-band activity throughout the hippocampal formation (Fig. S6). This rather subtle phenotype may still be of physiopathological relevance for disorders associated with KCC2 downregulation. Dentate spikes coincide with periods of stronger gamma-band activity coherence between hippocampus and cortex and are temporally correlated to the power of sharp-wave ripples^77^. Since both dentate spikes^78^ and sharp-wave ripples^79^ have been associated with memory consolidation in hippocampal-dependent learning, we predict KCC2 suppression in the pathology may impact cognitive performances through a combination of GABA-dependent and -independent mechanisms. Our results suggest therapeutic strategies aiming to either restore neuronal chloride homeostasis by blocking NKCC1 function^57^ and neuronal excitability by targeting leak potassium channels^80^, or to stabilize KCC2 membrane expression^25,32^ may best compensate for altered network activity and associated behavioral or cognitive deficits.

## Methods

### Animals

Juvenile male Wistar rats were obtained from Janvier Labs (Le Genest Saint Isle, France). All procedures conformed to the International Guidelines on the ethical use of animals, the French Agriculture and Forestry Ministry guidelines for handling animals (decree 87849, licence A 75-05-22) and were approved by the Charles Darwin ethical committee (APAFIS2015111011588776v3).

### Virus and constructs

Rat *slc12a5*-specific and non-target shRNA sequences^33^ were introduced in pTrip vector under U6 promoter and used to produce purified lentiviral particles (titer 7-9 × 10^9^ TU/ml, UNC Vector Core Facility)^35^. For some experiments, these same sequences were introduced in pAAV vector under U6 promoter and used to produce purified AAV particles (AAV2.1-shRNA-GFP, titer 10^13^ TU/ml, Atlantic Gene Therapy, Nantes). AAV particles expressing inhibitory DREADD under synapsin promoter were obtained from the Viral Vector Facility of Zurich University (AAV2.1-hM4D(Gi)-mCherry, titer 10^13^ TU/ml).

### Stereotaxic viral injection and EcoG electrodes implantation

30-days Wistar rats were anesthetized by intraperitoneal injection of ketamine / xylazine (75/10 mg/kg) and subcutaneous injection of buprenorphine (0.05 mg/kg) for analgesia. Animals were then head-fixed in a stereotaxic apparatus. 1 μl of lentivirus (concentrated at 10^9^ TU/ml) was injected bilaterally in the dentate gyrus (AP −4.0 mm, ML +/− 2.5 mm, DV −3.1 mm) at 125 nl/min. For DREADD experiments, 500nL of mix AAV2.1-shRNA-GFP and AAV2.1-hM4D(Gi)-mCherry (ratio 2/3:1/3) was injected bilaterally in the dentate gyrus. For experiments in CA1, animals were injected bilaterally with 1 μl of AAV2.1-shRNA-GFP (AP - 4.0 mm, ML +/− 2.5 mm, DV −2.35 mm).

For pilocarpine experiments, animals were also implanted with EcoG electrodes soldered to stainless steel screws. Two electrodes were placed above each hippocampus, at the same coordinates as the injection sites. A third screw was placed above the cerebellum and used as reference.

### Intracerebral probe implantation

For intra-hippocampal recordings, animals were anesthetized with isoflurane one week following viral injection, and injected subcutaneously with 0.05 mg/kg of buprenorphine for analgesia. One 16-channels silicon probe was implanted in the right hemisphere, at the site of the injection. Signals were monitored during the insertion of the probe to ensure correct localization of the probe in the hippocampus. One stainless steel screw was placed above the cerebellum and connected to the probe as a reference. Four additional screws were placed above the prefrontal cortex and the contralateral hemisphere to secure the dental cement adhesion. During the recovery period (minimum 3 days), rats were daily monitored and additional injection of buprenorphine (subcutaneous, 0.05 mg/kg) were made in case of detectable pain.

### Electrophysiology

#### Slice preparation

10 to 20 days after viral injection, animals were deeply anesthetized with ketamine / xylazine (115/15 mg/kg) and transcardially perfused with an ice-cold choline-based solution containing (in mM) : 110 Choline Cl, 25 Glucose, 25 NaHCO_3_, 11.6 Ascorbic acid, 3.1 Pyruvic acid, 1.25 NaH_2_PO_4_, 2.5 KCl, 0.5 CaCl_2_, 7 MgCl_2_ saturated with 95% O_2_/5% CO_2_. Rats were then decapitated, hippocampi were rapidly dissected and 400μm transverse sections were prepared using a vibratome (Microm, Thermofisher). Slices were then transferred and allowed to recover for 1 hour in a humidified interface chamber filled with bicarbonate-buffered ACSF pre-heated at 37°C and oxygenated with 5% CO_2_ in O_2_, containing (in mM) : 126 NaCl, 26 NaHCO_3_, 10 Glucose, 3.5 KCl, 1.25 NaH_2_PO_4_, 1.6 CaCl_2_, 1.2 MgCl_2_. For recordings, slices were transferred in a submerged recording chamber and superfused with ACSF maintained at 32°C.

#### Patch clamp recordings

Electrophysiological recordings were made with 4-5 MΩ borosilicate glass pipettes. Signals were acquired with a Multiclamp 700B amplifier (Molecular Devices), low-pass filtered at 10 kHz, and digitized at 20 kHz. All cells were recorded in the suprapyramidal blade of the dentate gyrus.

For perforated-patch recordings, internal solution contained (in mM) 140 KCl and 10 HEPES (pH adjusted to 7.4 with KOH). Gramicidin was added in the pipette solution to reach a concentration of 60 μg/ml. Rubi-GABA was added in ACSF (15 μM, Tocris) together with NBQX (20 μM, HelloBio), APV (50 μM, HelloBio), CGP54626 (10 μM, Tocris) and TTX (1 μM, Latoxan). Rubi-GABA was photolyzed using a digital modulated diode laser beam at 405 nm (Omicron Deepstar, Photon Lines, Marly Le Roi, France) delivered through a single path photolysis head (Prairie Technologies, Middleton, USA). The laser beam diameter was set to a diameter of 3-5 μm and was directed onto the soma of the recorded neuron. Photolysis was induced by a 5 ms/30 mW pulse. Once access resistance was stabilized, cells were held at a potential of −70 mV and 3.5 s voltage steps ranging from −100 to −30 mV were applied to the cell. Laser pulses were delivered at 2.2 s after the onset of the voltage step to allow for stabilization of the holding current. Currents were analyzed offline using Clampfit software (Molecular Devices). Voltages were corrected for liquid junction potential (4.1 mV) and for the voltage drop through the series resistance

For all other patch clamp experiments, cells were recorded in whole-cell configuration and maintained at −70 mV. All voltages were corrected offline for liquid junction potential (of 14.5 mV).

Intrinsic properties were evaluated using an internal solution containing (in mM) 120 K-Gluconate, 10 KCl, 10 HEPES, 0.1 EGTA, 4 MgATP, 0.4 Na_3_GTP (pH adjusted to 7.4 with KOH) and blocking synaptic transmission with NBQX, APV and picrotoxin (10μM, HelloBio). Resting membrane potential was measured using I=0 current clamp mode one minute after break-in. Input resistance was measured in current-clamp mode with −50 pA current steps. Input/output curves were derived from series of 800 ms current steps (ranging from −100 to 375 pA, 25 pA step increment). For potassium currents measurements, TTX (1 μM) was added to the bath. Currents were recorded in voltage-clamp mode between −120 mV and −60 mV with a continuous ramp at 0.03 mV/ms. For EPSP/spike coupling recordings, artificial waveforms derived from mEPSC recordings were used as a current command applied to the patch pipette. Effect of CNO application on membrane potential was estimated by whole-cell recordings in current-clamp mode with no current injection.

For miniature excitatory postsynaptic current (mEPSCs) recordings, pipettes were filled with internal solution containing (in mM) 115 Cs-Methylsulfonate, 20 CsCl, 10 HEPES, 0.1 EGTA, 4 MgATP, 0.4 Na_3_GTP (pH adjusted to 7.4 with CsOH) and currents were isolated by adding TTX and bicuculline to the extracellular solution. For miniature inhibitory postsynaptic currents (mIPSCs), pipettes were filled with (in mM) 135 CsCl, 10 HEPES, 10 EGTA, 4 MgATP, 0.4 Na_3_GTP, 1.8 MgCl_2_ (pH adjusted to 7.4 with CsOH) and currents were isolated by adding TTX, NBQX and AP-V in the bath. Series and input resistance were regularly monitored with −5 mV voltage steps and recordings were interrupted if either value varied by more than 20 Miniature synaptic currents were detected and analyzed offline using Detectivent software^81^.

#### Extracellular field recordings

For extracellular recordings, a recording pipette of 2-3 MΩ resistance was filled with ACSF and inserted within the granular layer, in a densely infected area of the dentate gyrus. A tungsten bipolar electrode (0.5 MΩ) was used to stimulate perforant path inputs. Stimulation intensity was adjusted to induce a population spike of about half its maximum amplitude. Field EPSP (fEPSP) slope was determined over a 1ms window preceding the population spike. Population spike amplitude was assessed as the distance between its peak and the baseline extrapolated from its initiation and termination points using a routine written under Matlab (The MathWorks, Inc., Natick, MA, USA).

### Biochemistry

#### Co-immunoprecipitation assays from rat hippocampal homogenates

Hippocampi were dissected from adult Sprague Dawley female rats and homogenized by sonication in co-immunoprecipitation buffer as above (1 ml/100 mg of tissue) and solubilized by rotation for 2 to 4 hours at 4°C. After centrifugation at 20,000 g for 40 minutes at 4°C, protein content of the supernatant was measured by BCA protein assay (Pierce). 6.5 mg of proteins were transferred into microtube and 5 μg of primary antibody (Rabbit anti-KCC2, Millipore, and Rabbit anti-Task3, Alomone) or nonimmune IgG control antibody of the same species (Jackson Immunoresearch) were added. Samples were then incubated overnight at 4°C under rotation and processed as above.

#### Co-immunoprecipitation assays from HEK293T cells

HEK293T cells were grown in DMEM GlutaMAX (Invitrogen) supplemented with 5 g/l glucose and 10% fetal bovine serum. Cells (60-70% confluent) were co-transfected using transfectin (Biorad) according to the manufacturer’s instructions with plasmids expressing Flag-tagged rat KCC2 (Chamma et al., 2013) and HA-tagged mouse Task3 (kindly provided by Guillaume Sandoz, Nice Univ., France) or HA-tagged GFP (Addgene) as control (with a 1:1 ratio). 48 hours after transfection, cells were homogenized by sonication in co-immunoprecipitation buffer containing (in mM): 50 Tris-HCl pH 7.4, 150 NaCl, 1 EDTA, as well as 0.5 % triton X-100 and protease inhibitors. Cells were then solubilized by rotation for 2 hours at 4°C and centrifuged at 20,000 g for 30 minutes at 4°C. Supernatants were transferred into a microtube with 20 μl of pre washed anti-HA coupled beads (Cell signaling) or EZview™ Red anti-Flag M2 Affinity Gel (Sigma). Samples were incubated overnight at 4°C under rotation. Complexes were precipitated with 20 μl protein-G magnetic beads (Invitrogen) for 2 hours at 4°C. Beads were then washed twice with 1ml of co-immunoprecipitation buffer and once with triton-free co-immunoprecipitation buffer. Bound complexes were eluted in 4X sample buffer (Invitrogen) for 1 hour at 37°C. Samples were analyzed by SDS-PAGE and western blot.

#### Western blotting

Proteins were separated on a 4-12% SDS polyacrylamide gradient gel (Invitrogen) and transferred onto a nitrocellulose membrane (GE Healthcare). For co-immunoprecipitation assays from rat hippocampal homogenate, blots were probed with antibodies against KCC2 (1/1000, Millipore) and Task3 (1/250, Santa Cruz). For co-immunoprecipitation assays from HEK293T cells, blots were probed with HA (1/3000, Cell signaling) and KCC2 or FlagM2 (1/1000, Sigma). The primary antibodies were detected with fluorescent secondary antibody (1/1000-1/3000, DyLight 700 or 800, Rockland) using Odyssey infrared imaging system (LI-COR Bioscience). All biochemical assays were repeated at least 3 times on independent hippocampal extracts or cultures.

### Immunocytochemistry

Neuro-2a cells were grown on glass coverslips in DMEM GlutaMAX (Invitrogen) supplemented with 1 g/l glucose and 10% fetal bovine serum. Cells at 60-70% confluence were co-transfected using Transfectin (Biorad) according to manufacturer’s instructions with plasmids expressing HA-tagged mouse Task-3 (kindly provided by Guillaume Sandoz, Nice Sophia Antipolis Univ., France) and either Flag-tagged rat KCC2^7,11^ or a pCAG empty vector as control (with a 1:3 ratio). 48 hours after transfection, cells were washed with PBS, fixed with paraformaldehyde (4%) supplemented with 4% sucrose in PBS and permeabilized with 0.25% Triton X-100 in PBS. After 3 × 10 min washes in PBS, cells were incubated for 30 min in blocking buffer (10% normal goat serum in PBS). Incubation with primary antibodies, HA (rabbit, 1/2000, Chemicon) and FlagM2 (mouse, 1/2000, Sigma) was then performed in blocking buffer for 2 hours at room temperature. After 3 × 10 min washes in PBS, cells were incubated with secondary antibodies for 1 hour (donkey anti-rabbit Alexa 647, 1/1000, Jackson Immunoresearch and goat anti-mouse CY3, 1/2000, Jackson Immunoresearch). Coverslips were washed 3 times in PBS and mounted in Mowiol/Dabco (25 mg/ml) solution. Immunofluorescence images were acquired using an upright confocal microscope (Leica TCS SP5), using a 63× 1.40-N.A. objective with 2.5X electronic magnification and an Ar/Kr laser set at 561 and 633 nm for excitation of Cy3 and Alexa 647, respectively.

### Immunohistochemistry

At least 2 weeks following viral injections, rats were anesthetized by intraperitoneal ketamine / xylazine injection (110/15 mg/kg) and perfused transcardially with ice-cold cutting solution of choline as above. Brain were removed, postfixed in PFA 4% for 48 hours and stored at 4°C in 30% sucrose PBS. Parasagittal sections (40 μm thick) were obtained using a cryotome. After washes, brain slices were preincubated 3 hours with 0.5% Triton and 10% goat serum in PBS and then 48 hours at 4°C with primary antibodies: GFP (chicken, 1:1000, Millipore), KCC2 (rabbit, 1:400, Millipore), Prox1 (rabbit, 1:10000, Millipore), Task3 (rabbit, 1:400, Alomone), Trek2 (rabbit, 1:400, Alomone). After rinsing in PBS, slices were incubated 3 hours with secondary antibody (Cy3-coupled goat anti-rabbit and FITC-coupled goat anti-chicken) and mounted with Mowiol/Dabco (25 mg/mL). Images were acquired on an upright confocal microscope (Leica TCS SP5), using a 63× 1.40-N.A. objective with 2X electronic magnification and Ar/Kr laser set at 491 and 561 nm for excitation of Cy3 and FITC, respectively. Stacks of 4 to 6 μm optical sections were acquired at a pixel resolution of 1024 × 1024 with a z-step of 0.2 μm. Individual sections were then analyzed with Imaris (Bitplane). For images of the entire dentate gyrus, images were acquired using a 10X objective and stacks of 10 to 20 μm optical sections were acquired at a pixel resolution of 1024 × 1024 with a z-step of 1 μm.

### Behavior and recordings

All recordings took place in a dimly-lit area enclosed by black curtains. Animals were handled for at least 3 days for habituation and to ensure stability of the recordings (location of the probe, power of the signal) before experiments started. A 80 cm × 80 cm arena with 50 cm high black plastic walls was used for exploration phase. A white card on one of the wall was used as spatial cue. For sleep recordings, animals were put in a white cylinder box (20 cm of diameter, 40 cm high). During the habituation phase, animals were placed in the cylinder box then in the empty arena for 10 min each, twice a day.

For recordings of awake behavior, new objects were placed in the open-field to stimulate exploration. All sleep recording sessions took place during the 2 hours following exploration of novel objects. For DREADD experiments, saline or CNO (1 mg/kg) was injected in i.p. 30 minutes before starting recording. All recordings took place less than 3 hours after i.p. injection.

Data were acquired at 20 kHz using an Intan recording controller (Intan Technologies, Los Angeles, USA) and Intan Recording Controller software (version 2.05).

### Data analysis

All analyses were performed offline using Matlab built-in functions and custom-written scripts as well as Chronux (http://chronux.org/) and the FMAToolbox (http://fmatoolbox.sourceforge.net/). Power spectra and spectrograms were computed using multi-tapers estimates on the raw LFP. Theta power was determined in the 5–10 Hz band. Dentate spikes detection was performed after high-pass filtering (> 30Hz), squaring and normalizing the field potential in the dentate gyrus. DS were defined as events of positive deflection peaking at > 8 s.d. and remaining above 4 s.d. for less than 30 ms. Current source density analysis (CSD) revealed two distinct DS profiles, previously described as DS1 and DS2^55^. Since most (70-80 %) DS in our recording conditions were of type 2, we did not further attempt to distinguish between DS subtypes in our analysis.

Ripple detection was performed by band-pass filtering (100–600 Hz), squaring and normalizing, followed by thresholding of the field potential recorded in CA1 pyramidal layer. SPW-Rs were defined as events starting at 4 s.d., peaking at >6 s.d., and remaining at >4 s.d. for <150 ms and >15 ms

### Electrocorticogram (EcoG) recordings and epilepsy induction

EcoG data were acquired at 20 kHz using Epoch Wireless recording system (Ripple, Salt Lake City, USA) and Clampex software (Molecular Devices). Video recording (29 frames/s) was synchronized to data acquisition to enable offline analysis.

For epilepsy induction, animals received intraperitoneal injection of 127 mg/kg of lithium 24 hours before treatment. The following day, animals received i.p. injection of 1 mg/kg of methylscopolamine in order to prevent peripheral effects of pilocarpine. 30 min later, 40 mg/kg of pilocarpine was injected i.p. Both video and EcoG data were acquired for one hour following pilocarpine injection. Status epilepticus was then interrupted by i.p. injection of 5 mg/kg diazepam. Animals were then sacrificed and brain were recovered and processed for visualization of the infected area.

Rat behavior was scored according to the following modified Racine’s scale : stage 1, immobility and chewing; stage 2, neck and body clonus; stage 3, forelimb clonus and rearing; stage 4, *status epilepticus*, uninterrupted seizures. The onset of *status epilepticus* was assessed based on EcoG recordings.

### Statistics

All statistical tests were performed using Matlab functions (Statistics and Machine Learning Toolbox). Data are presented as mean ± sem unless stated otherwise. All tests are two-tailed tests. Comparison of means was performed using Student’s t-test for normally distributed variables (as tested with Shapiro-Wilk test) of equal variances (tested with Bartlett test).

Otherwise, comparison of mean was performed using the non-parametric Mann-Whitney test. Kolmogorov-Smirnov test was used for comparison of distributions. Significance was determined as p < 0.05.

## Acknowledgements

We are grateful to Quentin Chevy for sharing original observations related to the present work. We also thank Richard Miles and Kai Kaila for critical reading of the manuscript. This work was supported in part by Inserm, Sorbonne Université, as well as the Fondation pour la Recherche Médicale (Equipe FRM DEQ20140329539 to J.C.P.), the Human Frontier Science Program (RGP0022/2013 to J.C.P. and L.M.P.), ERANET-Neuron (funded by ANR to J.C.P. and MINECO to L.M.P.) and the Fondation Française pour la Recherche sur l’Epilepsie - Fédération pour la Recherche sur le Cerveau (research grant to J.C.P.). M.G. was the recipient of fellowships from Sorbonne Université and Fondation pour la Recherche Médicale as well as an IBRO-InEurope Short Stay Grant. The Poncer lab is affiliated with the Paris School of Neuroscience (ENP) and the Bio-Psy Laboratory of excellence.

## Author contributions

M.G., L.M.P. and J.C.P. designed the research. L.M.P. and J.C.P. supervised the research. M.G. performed all experiments involving animal surgery, *in vitro* and *in vivo* electrophysiology, immunohistochemistry and imaging. S.A.A. performed all biochemical assays, immunocytochemistry and imaging of heterologous cells. E.F. and J.C.P. performed experimental epilepsy assays. D.G.D. and L.M.P. helped set up *in vivo* electrophysiological recordings and data analysis, T.I. contributed confocal imaging and quantification. M.G., S.A.A. and E.F. analyzed the data. M.G. and J.C.P. wrote the paper.

---

**Figure S1.**
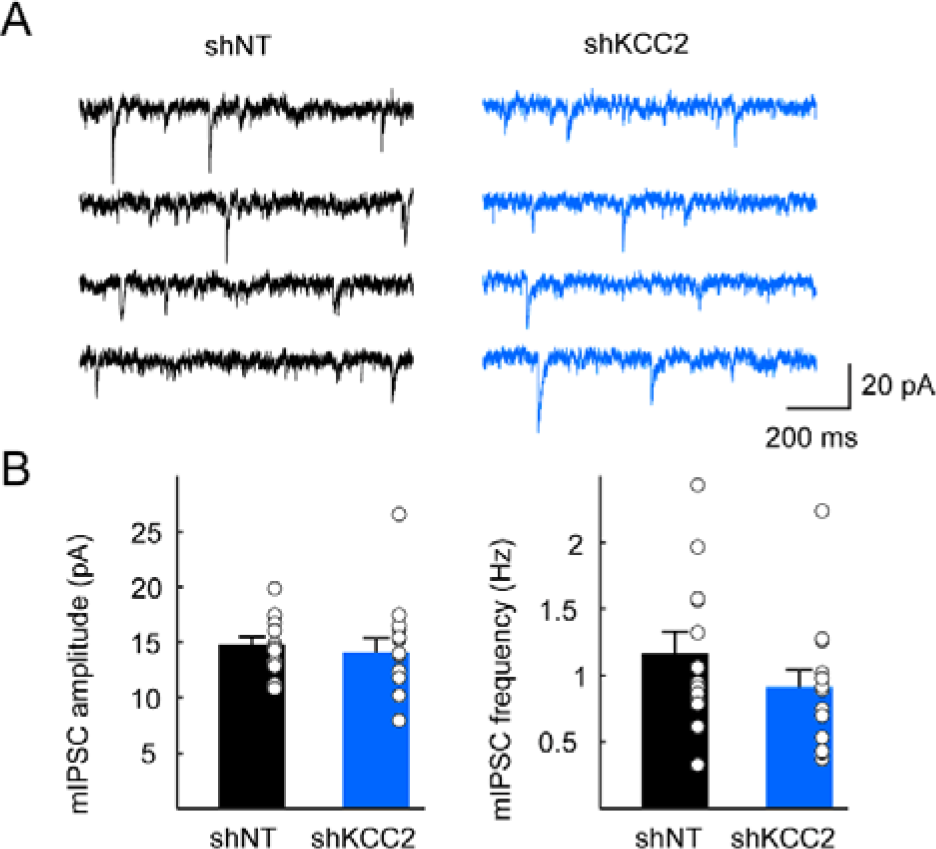
KCC2 suppression does not affect the efficacy of GABAergic synapses. **A**, 4 s-recordings of mIPSCs from granule cells expressing non-target (shNT, black) or KCC2-directed (shKCC2, blue) shRNA. **B**, Summary data from shNT (n = 13 cells, 2 rats) and shKCC2 (n = 14 cells, 2 rats) showing no changes in mIPSCs amplitude (left, p=0.42) or frequency (right, p=0.27).

**Figure S2.**
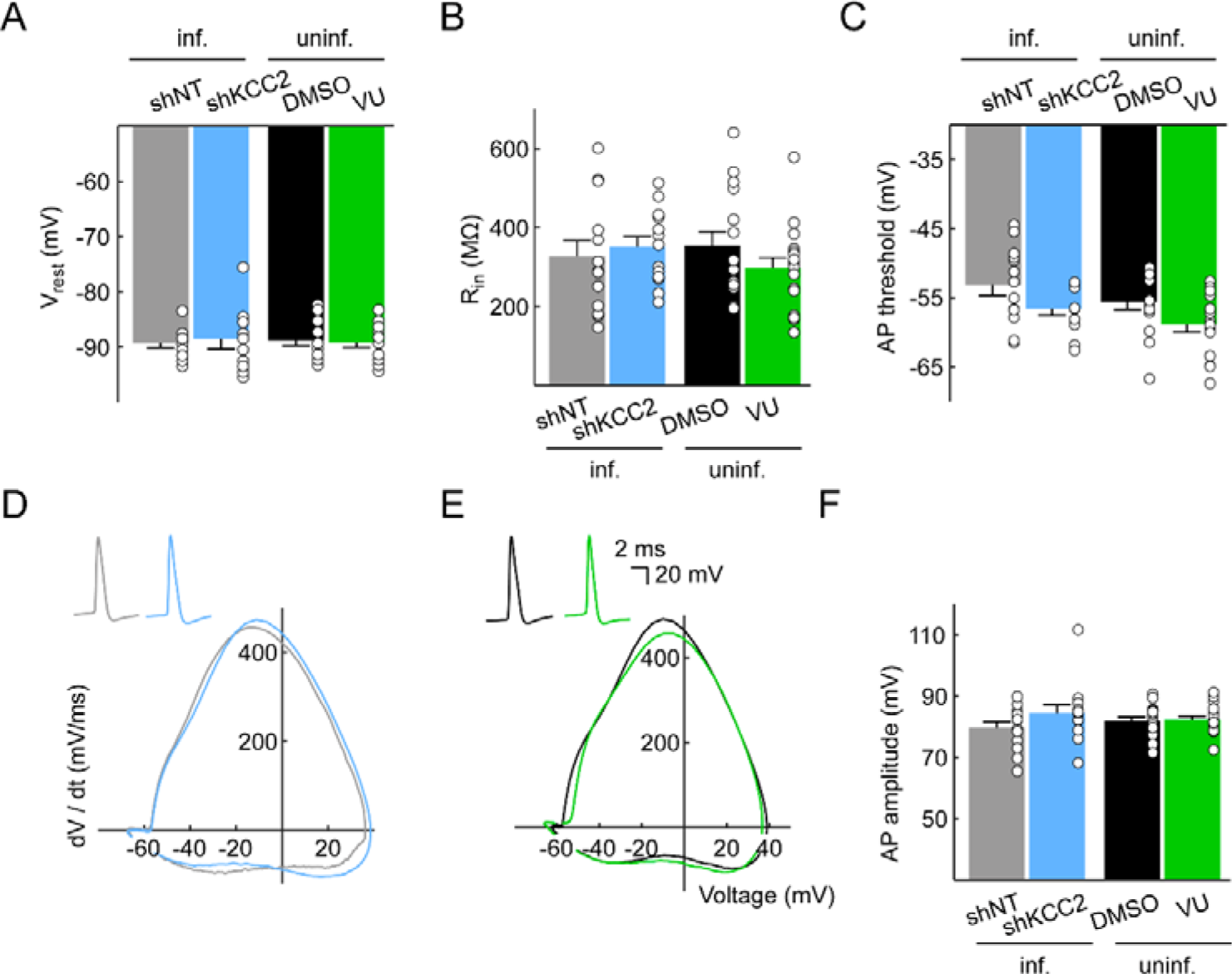
Increased excitability upon KCC2 knockdown is cell-autonomous and independent of KCC2-mediated ion transport. Membrane and spike properties were analyzed for uninfected cells from slices infected with lentiviral vectors expressing non-target (grey, n = 13 cells, 5 rats) or KCC2-directed (light blue, n = 13 cells, 7 rats) shRNA sequences. Effect of KCC2 blockade for >30 min with its specific antagonist VU0463271 was also analyzed (green, n = 17 cells, 5 rats) and compared to DMSO treatment (black, n = 15 cells, 4 rats). No differences were observed in membrane potential (**A**, shNT *vs* shKCC2 p = 0.83, DMSO *vs* VU p =0.75), input resistance (**B**, shNT *vs* shKCC2 p = 0.63, DMSO *vs* VU p =0.20) was well as AP threshold (**C**, shNT *vs* shKCC2 p = 0.063, DMSO *vs* VU p =0.055), waveform (**D-E**) and amplitude (**F**, shNT *vs* shKCC2 p = 0.24, DMSO *vs* VU p =0.77).

**Figure S3.**
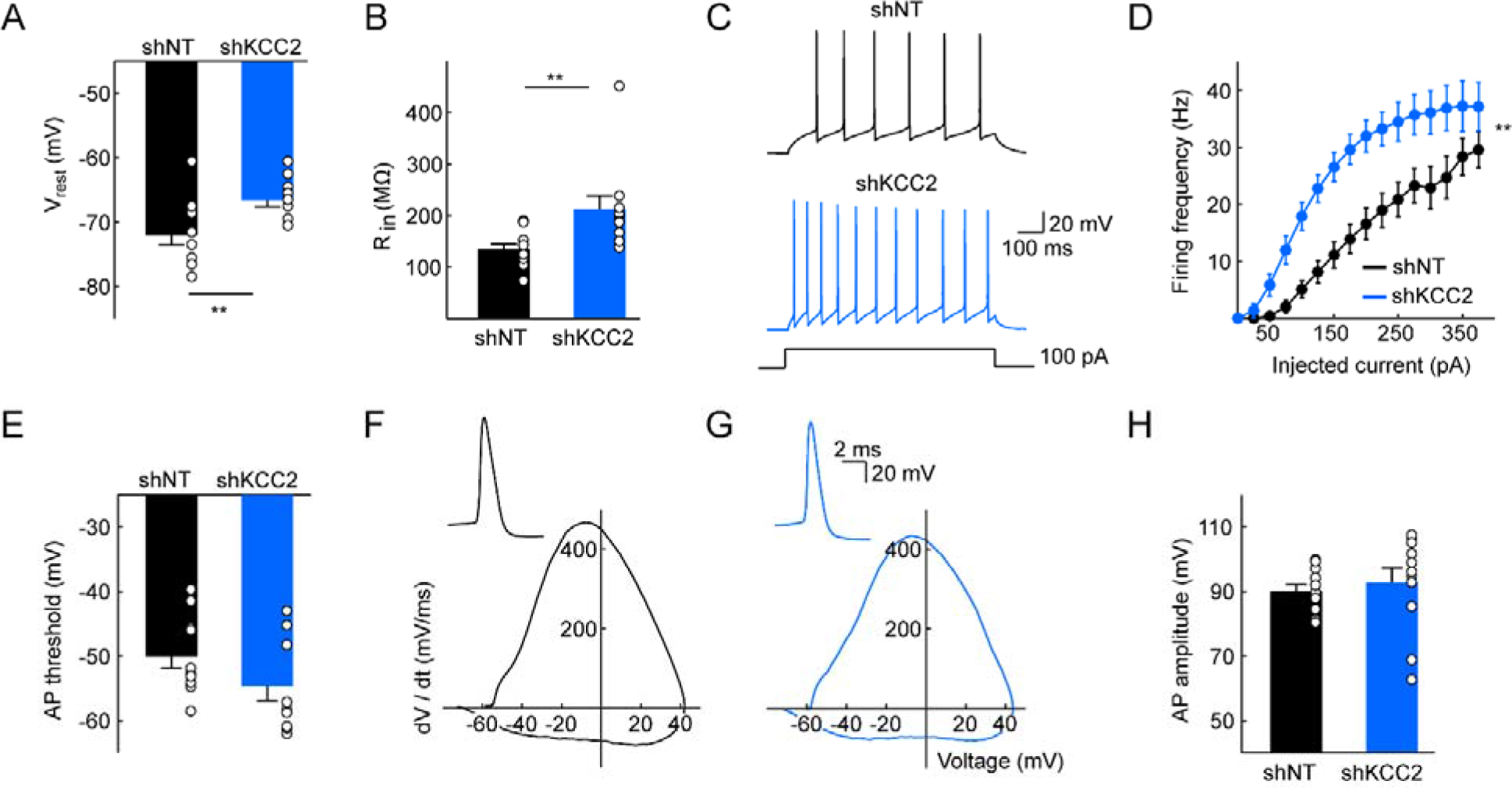
Increased excitability in CA1 pyramidal cells upon KCC2 suppression. **A-B**, Membrane properties of CA1 superficial pyramidal cells expressing non-target (n = 11 cells, 3 rats) or KCC2-directed (n = 11 cells, 3 rats) shRNA, as detected in whole-cell patch clamp recordings while blocking synaptic transmission. KCC2 knockdown resulted in depolarized membrane potential (A, ** p<0.01) and increased input resistance (B, ** p<0.01). **C**, Individual traces in both conditions for a depolarizing current pulse of 100 pA. **D**, Mean input/output relationships representing the frequency of APs as a function of injected current (** p<0.01). No difference was observed between shNT and shKCC2-expressing CA1 pyramidal cells with regard to action potential threshold (**D**, p=0.13), spike waveform (**E**) or spike amplitude (**F**, p=0.59).

**Figure S4.**
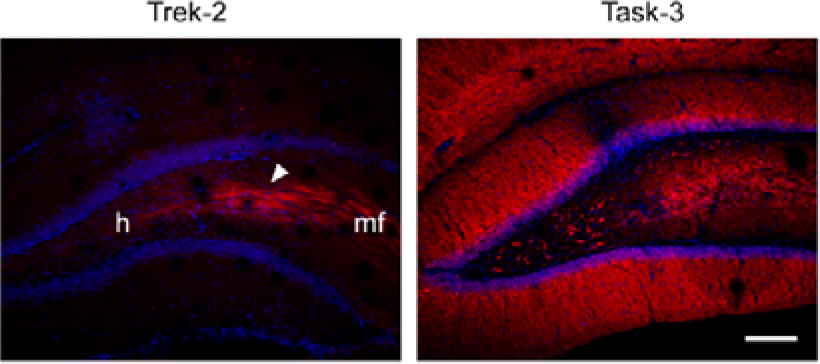
Differential subcellular localization of leak-potassium channels in the dentate gyrus. Left, Average intensity projection of confocal micrographs of dentate gyrus immunostained with Trek-2 antibody. Note the massive expression of Trek-2 in mossy fibers (arrowhead) and its relative absence in the granular and molecular layers of the dentate gyrus. h : hilus, mf : mossy fibers. Right, Average intensity projection of confocal micrographs of dentate gyrus immunostained with Task-3 antibody revealing a strong expression in both granular and molecular layers as well as in hilar mossy cells. Scale, 200 μm.

**Figure S5.**
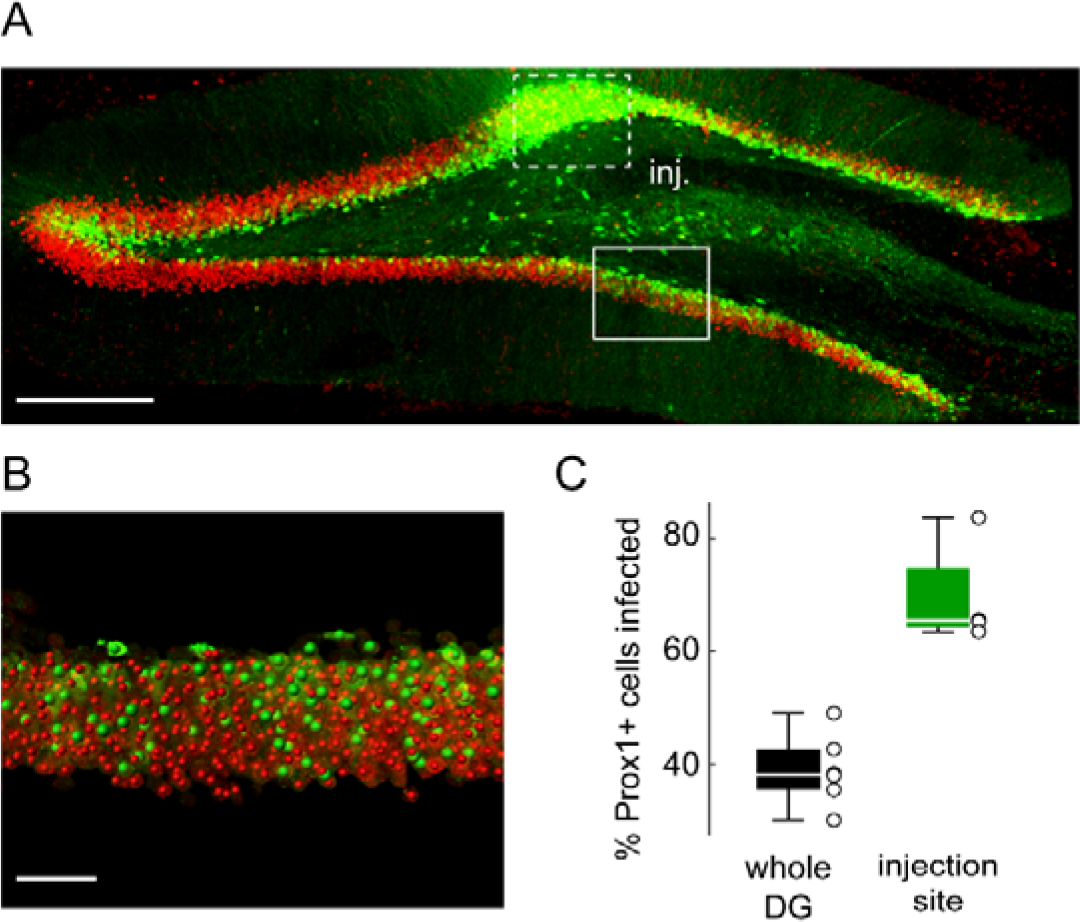
Lentiviral infection yield in the dorsal dentate gyrus. **A**, Maximal intensity projection of confocal micrographs showing infected cells (GFP, green) and Prox1 (red) immunostaining in a parasagittal hippocampal slice 2 weeks after lentivirus injection. Dotted box: injection site. Scale, 300 μm. **B**,Imaris 3D rendering of plain boxed area shown in A. For colocalization estimates, dentate granule cell nuclei were automatically detected based on Prox1 immunostaining (red spheres) while infected cells were detected based on GFP fluorescence (green spheres). **C**, Summary of mean infection rate through the whole dentate gyrus (n = 6 slices, 3 animals) and within 250 μm around the infection site (n = 4 slices, 2 animals).

**Figure S6.**
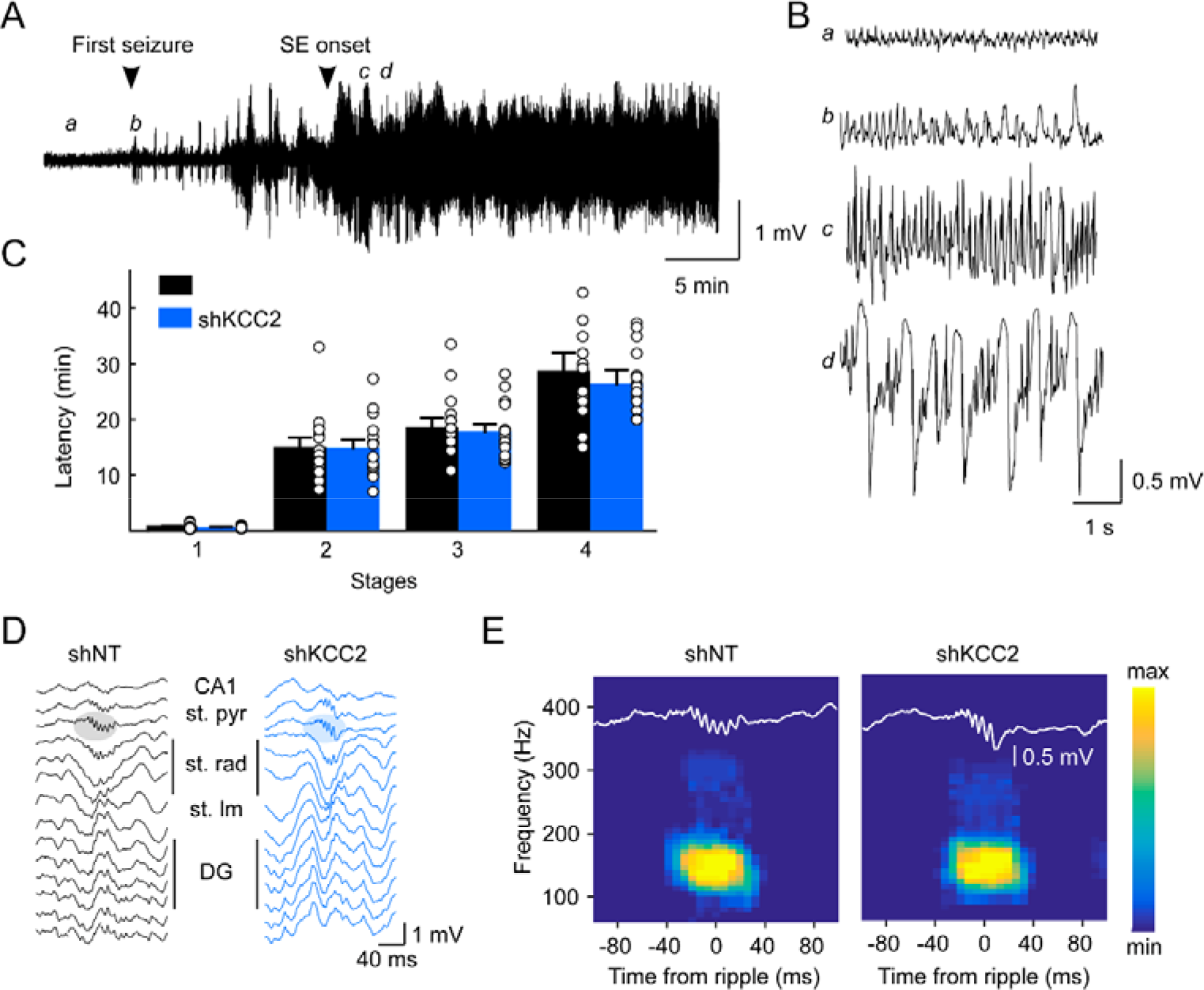
KCC2 suppression in the dentate gyrus does not induce or promote epileptiform activity. **A**, Example trace of an EcoG recording during epilepsy induction with a pilocarpine protocol. Arrowheads indicate the timing of the first seizure and the onset of the *status epilepticus*. **B**, Expanded sections of the recording shown in A, showing the evolution of the EcoG signal at different time points of ictogenesis. **C**, Behavioral state was classified according to modified Racine’s scale as follows: stage 1, immobility and chewing; stage 2, neck and body clonus; stage 3, forelimb clonus and rearing; stage 4, *status epilepticus*, uninterrupted seizures. The onset of *status epilepticus* was assessed based on EcoG recordings. Latency to different stages was similar between groups (n = 15 animals each). **D**, Traces of ripple events recorded in CA1 during sleep for rats implanted with 16-channels silicon probes as in **Fig. 5**. **E**, Ripples were automatically detected and 100 events per animal were randomly selected (shNT n= 5 rats, shKCC2 n = 7 rats). Suppression of KCC2 in the dentate gyrus did not lead to pathological fast ripples as evidenced by the absence of band around 300 Hz in the average time-frequency spectra.

**Figure S7.**
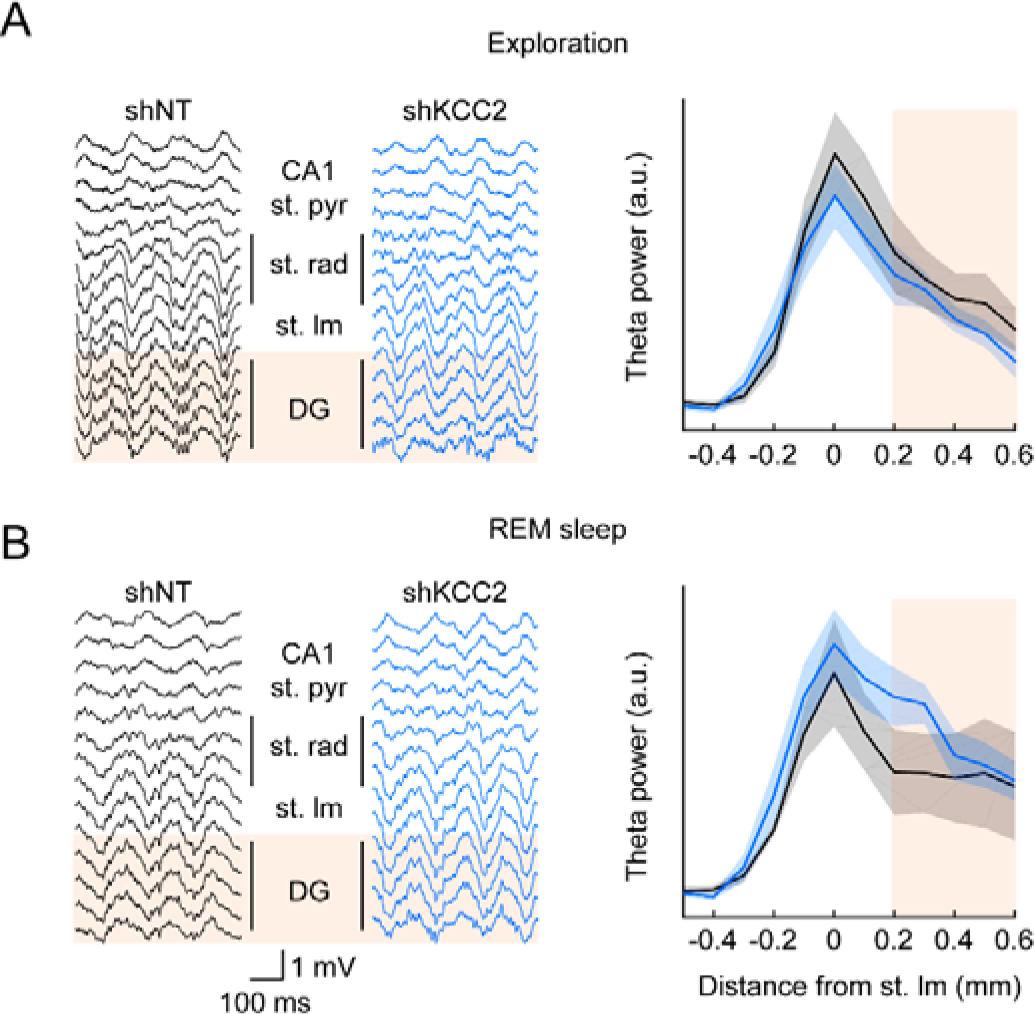
KCC2 suppression in the dentate gyrus does not affect hippocampal theta rhythm. **A**, Left, 500 ms stretches of intrahippocampal recordings (as in **Fig. 5**) during exploratory behavior in rats expressing non-target shRNA (left, black) or KCC2-directed shRNA (blue, right) in the dentate gyrus. Note the prominent theta oscillation in all channels. The localization of the *stratum lacunosum moleculare* was defined by the highest power of theta and CA1 was identified by the presence of action potentials and of ripples during sleep. Localization of the dentate gyrus is highlighted in orange. Right, theta power in the 5-10 Hz frequency band measured over > 1 min of recording using multi-tapers estimates. Power is represented as a function of the electrode localization. Depth of the electrode is expressed as the distance from *stratum lacunosum moleculare* (SLM). (2-way ANOVA, F_11,144_ = 2.03, p = 0.15. **B**, Similar as A, but during REM sleep. Depth profile was slightly different between groups (2-way ANOVA, F_11,133_ = 5.96, p = 0.016). However, comparisons for each layer individually did not yield any significant difference (Mann-Whitney, p > 0.05 for all layers).

**Figure S8.**
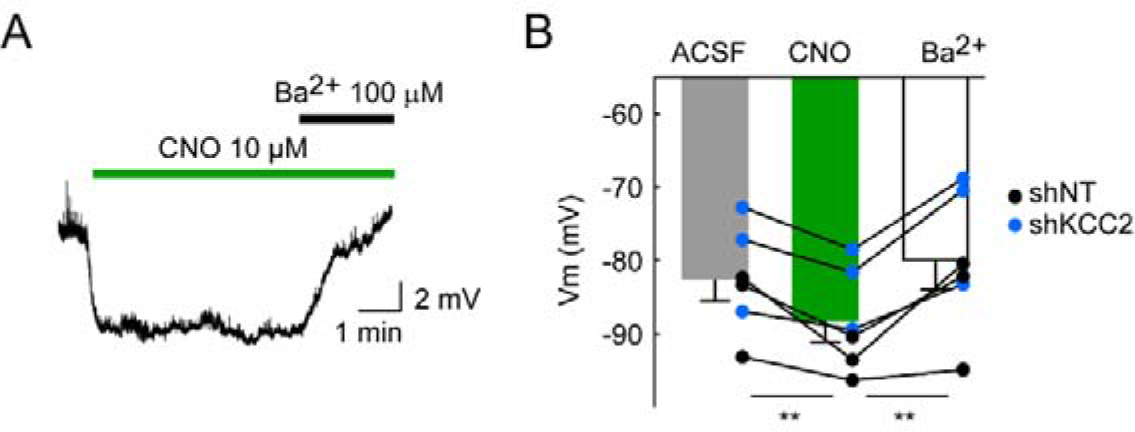
CNO application hyperpolarizes hM4D(Gi)-expressing dentate granule cells. **A**, Example current clamp recording of a dentate granule cell expressing non-target shRNA and hM4D(Gi) receptor, showing hyperpolarization of the membrane potential upon CNO (10 μM) bath application. Application of Ba^2+^ (100 mM BaCl_2_) completely blocks CNO-induced hyperpolarization. **B**, Summary data from 3 cells expressing non-target shRNA and 3 cells expressing KCC2-directed shRNA, together with hM4D(Gi). All cells were hyperpolarized following CNO application (** p<0.01) and repolarized upon Ba^2+^ application (** p<0.01).

